# Parvalbumin interneuron circuit sharpens orientation tuning via the extraclassical surround in cat V1

**DOI:** 10.64898/2026.11.13.766262

**Authors:** Andrzej T. Foik, Yongjun Liu, Huanjun Shao, Georgina Lean, Li Zhang, Leo R. Scholl, Roberto Jappelli, David C. Lyon

**Affiliations:** Department of Anatomy and Neurobiology, School of Medicine, University of California, Irvine, Irvine, CA, USA; International Centre for Translational Eye Research, Institute of Physical Chemistry, Polish Academy of Sciences, Warsaw, Poland; Institute of Physical Chemistry, Polish Academy of Sciences, Warsaw, Poland; State Key Laboratory of Resource Insects, Institute of Apicultural Research, Chinese Academy of Agricultural Sciences, Beijing, China; Laboratory of Genetics, The Salk Institute for Biological Studies, La Jolla, CA, USA

## Abstract

Parvalbumin-expressing (PV) inhibitory neurons are central to cortical gain control, yet optogenetic studies disagree on whether they sharpen orientation tuning or merely rescale responses. Because this work has been performed almost exclusively in rodents, which lack orientation columns and an orientation-matched extraclassical surround, the contribution of surround circuitry has remained untested. Here we combine cell-type-specific rabies-virus tracing with optogenetics to activate or suppress PV circuits in cat primary visual cortex, and measure single-neuron orientation tuning for stimuli confined to the classical receptive field or extending into the extraclassical surround. Activating PV circuits sharpened tuning progressively with illumination power for classical-receptive-field stimuli, but left tuning unchanged once the surround was engaged; conversely, suppressing them broadened tuning only for surround-engaging stimuli and left classical-receptive-field tuning unchanged. This double dissociation places PV circuitry in the causal path that sets tuning width and identifies surround suppression as the inhibition it supplies: where the surround is not recruited there is little PV-mediated inhibition to remove, and where it is, removing that inhibition widens tuning. PV-dependent sharpening is therefore bounded by the receptive-field architecture through which the population is recruited, not by a fixed property of the PV population itself.

## Introduction

A balance between excitation and inhibition underlies virtually every cortical computation, and inhibitory interneurons expressing the calcium-binding protein parvalbumin (PV) are among its principal regulators. In the primary visual cortex (V1), PV interneurons target the somata and proximal dendrites of excitatory pyramidal cells and, driven by local recurrent excitation, exert powerful control over local gain^1–3^. PV cells are themselves inhibited directly by somatostatin-expressing (SST) interneurons^4–6^ and indirectly through SST synapses onto the distal dendrites of pyramidal neurons^7,8^, so their output reflects the state of the surrounding inhibitory network rather than pyramidal drive alone.

Despite a decade of optogenetic work in mouse V1^9^, the role of PV interneurons in shaping orientation selectivity remains unresolved. Activating PV cells has been reported to rescale responses divisively without changing tuning width, and to sharpen tuning, with the outcome depending on activation strength and stimulus context ^1–3^. Several studies instead attribute sharpening to SST interneurons, whose suppressive effect is weaker^10^ but more selective^11,12^, and conclude that PV cells do not sharpen tuning at all^1,2^.

One explanation for this disagreement has not been testable in mice. Mouse V1 lacks orientation columns and the orientation-matched organization of the extraclassical surround found in carnivores and primates^13–15^. In columnar visual cortex, surround suppression is spatially structured and tightly coupled to orientation domains, and surround stimulation sharpens orientation tuning on its own^16–25^. If PV-mediated inhibition and surround suppression draw on overlapping circuitry, then whether PV activation sharpens tuning should depend on whether the surround is engaged, a prediction that requires a species with an orientation map to test.

Here we test that prediction in cat V1. We used a monosynaptic rabies-virus strategy^26–28^ to deliver channelrhodopsin-2 (ChR2) or archaerhodopsin-T (ArchT) to PV interneurons and to the neurons that synapse directly onto them, a configuration we refer to throughout as the PV-related circuit, and we compared it with a construct restricted to PV cells themselves. Recording single neurons in anaesthetized cats, we measured orientation tuning during bidirectional optogenetic manipulation, using stimuli sized either to the classical receptive field (CRF) of each neuron or extended to engage its extraclassical surround (full-field, FF). Activation sharpened and suppression broadened tuning for CRF stimuli, whereas activation no longer sharpened the already-narrow tuning measured with surround-engaging stimuli. PV-dependent sharpening is therefore not a fixed property of the PV population but is bounded by the receptive-field architecture through which the population is recruited.

## Results

### Cell-type-specific delivery of optogenetic actuators to PV circuits in cat V1

To gain optogenetic access to PV interneurons and their presynaptic partners in a species without transgenic tools, we built an adeno-associated viral (AAV) helper construct in which the previously characterized fugu PV promoter^29^ drives a polycistronic cassette encoding yellow fluorescent protein (Y), the avian TVA receptor (T) and rabies glycoprotein (B; RabG) (Fig. 1a). Expression of this cassette restricts infection by an EnvA-pseudotyped, glycoprotein-deleted rabies virus (RV) to TVA-expressing PV cells (Fig. 1b); complementation by the AAV-supplied RabG then allows the RV to spread retrogradely one synapse to label presynaptic partners (Fig. 1c). The RV carried either ChR2 with mCherry or ArchT with GFP, the fluorescent reporter serving for histological identification, which allowed us to activate or suppress the PV-related circuit while recording extracellularly with multichannel electrodes (Fig. 1d).

**Fig. 1.**
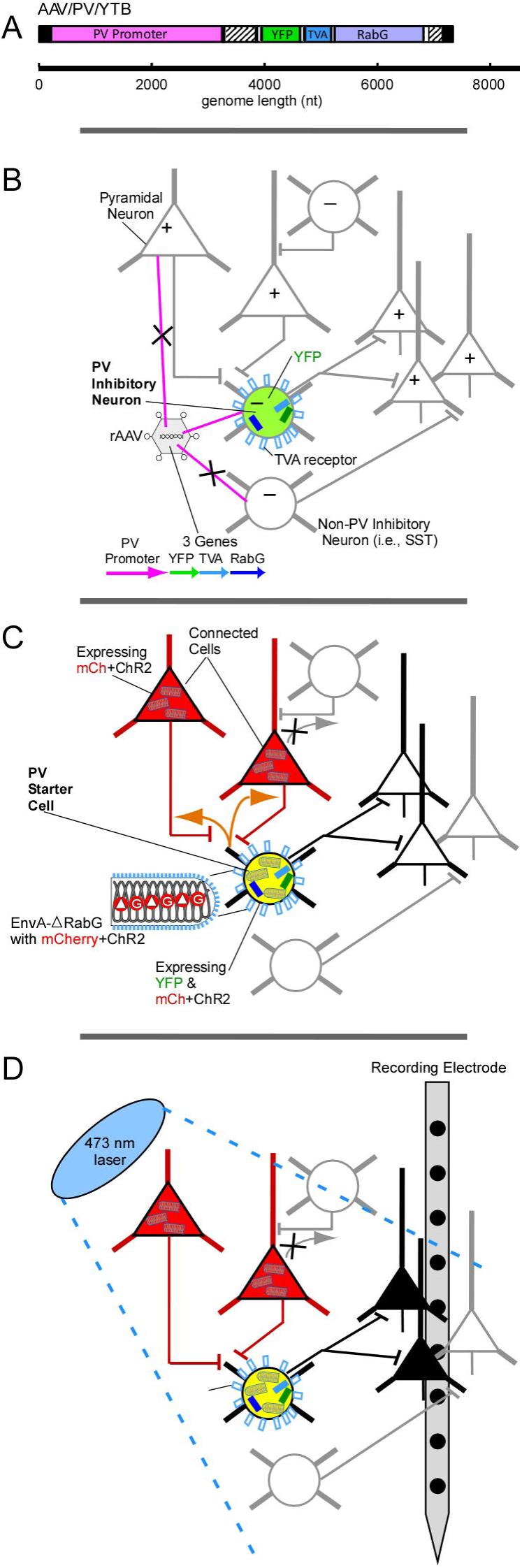
| Strategy for infecting the PV-related circuit. **a**, Genome map of the AAV-PV-YTB construct: the fugu PV promoter drives a polycistronic cassette encoding YFP, TVA and rabies glycoprotein (RabG). b, The recombinant AAV produces the TVA-YFP-RabG cassette only in parvalbumin-expressing interneurons, leaving pyramidal and non-PV inhibitory neurons uninfected. c, EnvA-pseudotyped, glycoprotein-deleted rabies virus carrying mCherry and ChR2 infects TVA-expressing PV starter cells and, following complementation with AAV-supplied RabG, spreads retrogradely one synapse to presynaptic partners. d, Configuration for optogenetic stimulation and multichannel recording in cat visual cortex.

Co-localization of PV immunofluorescence with the AAV-driven YFP reporter confirmed that the construct was selective for PV interneurons, and double labelling with the RV reporter identified starter cells (Fig. 2). Expression of the RV-delivered reporter was strong enough to visualize injection sites through the intact cortical surface in vivo, which allowed illumination and recording to be targeted to infected tissue (Fig. 3a–c), and post-mortem reconstruction confirmed electrode placement within the labelled network (Fig. 3d). Starter cells, identified by co-expression of the YFP and RV reporters, were parvalbumin-immunopositive in the great majority of cases, and none showed pyramidal morphology (Fig. 2).

**Fig. 2.**
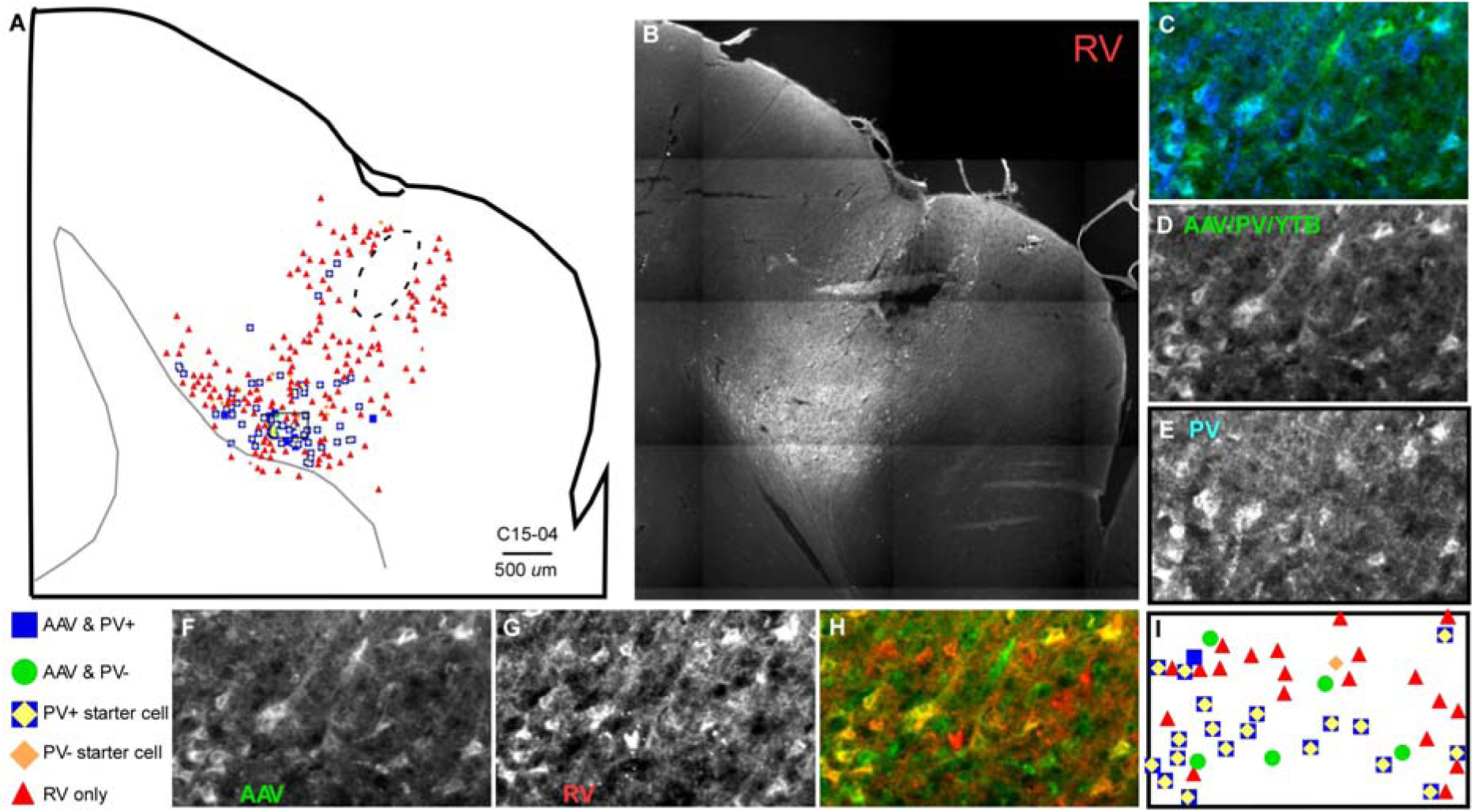
| Anatomical verification of PV-related circuit tracing. **a**, Reconstruction of the distribution of labelled cells relative to the AAV/RV injection site (case C15-04), showing double-labelled starter cells (yellow diamonds, PV⁺; orange diamonds, PV⁻), AAV-only PV⁺ cells (blue squares), AAV-only PV⁻ cells (green circles) and RV-only presynaptic cells (red triangles). Scale bar, 500 µm. b, Low-power fluorescence image of the rabies virus (mCherry) injection site corresponding to a. c–e, High-power images of the same field: c, AAV-PV-YTB (YFP, green) combined with parvalbumin immunofluorescence (blue); d, cells expressing the TVA-YFP-RabG cassette; e, parvalbumin immunolabelling. f–h, High-power images of AAV and rabies virus double-labelled sections: f, AAV-infected parvalbumin-expressing cells (YFP, green); g, rabies-infected PV-related circuit (mCherry, red); h, merged fluorescence showing co-infected starter cells (yellow). i, Map of cell types in the reconstructed field, using the symbols and colours of a.

**Fig. 3.**
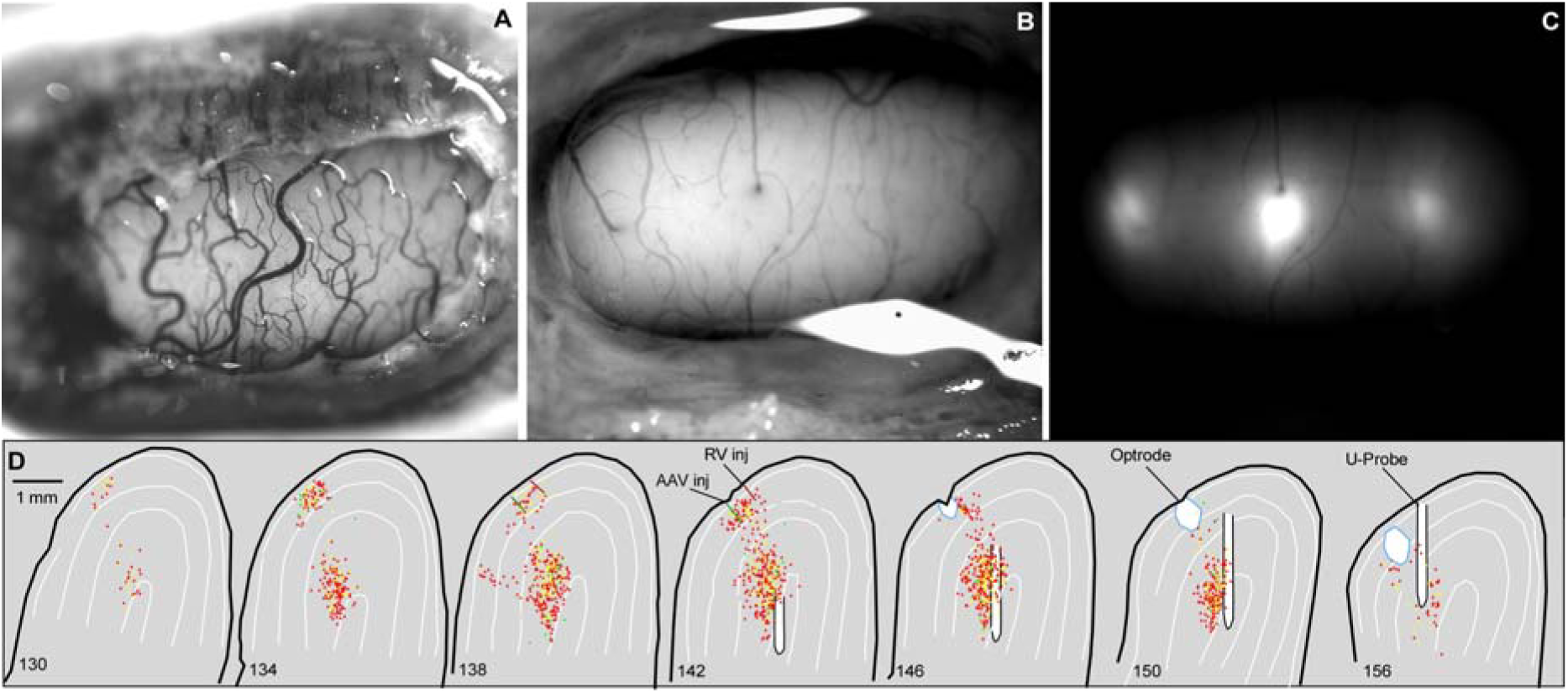
| Rabies-driven fluorescence in the PV-related circuit is visible in vivo. **a**, Freshly exposed surface of cat V1. b, The same surface after EnvA rabies virus injection at three locations. c, The same region immediately before the optogenetic experiment, showing mCherry fluorescence at the three injection sites through the intact cortical surface. d, Anatomical reconstruction of the injection site at which optogenetic stimulation and single-neuron recording were performed. Yellow diamonds, starter cells co-infected by AAV and rabies virus; red triangles, presynaptic cells labelled by rabies spread.

### Bidirectional optogenetic control of firing rate in the PV-related circuit

We first asked whether the two actuators controlled activity in the expected directions. In a minority of recorded neurons, blue illumination of ChR2-expressing tissue increased firing both in the absence of visual stimulation and during drifting gratings (Fig. 4a–d; Supplementary Fig. 1a–d); these cells are presumed to express the actuator directly. In the majority, the same illumination reduced visually driven firing (Fig. 4e–g; Supplementary Fig. 1e–i), as expected for neurons receiving enhanced PV-mediated inhibition. Yellow illumination of ArchT-expressing tissue produced the complementary pattern, suppressing firing in presumed actuator-expressing cells (Fig. 4j–m) and increasing it in the wider population (Fig. 4n–p).

**Fig. 4.**
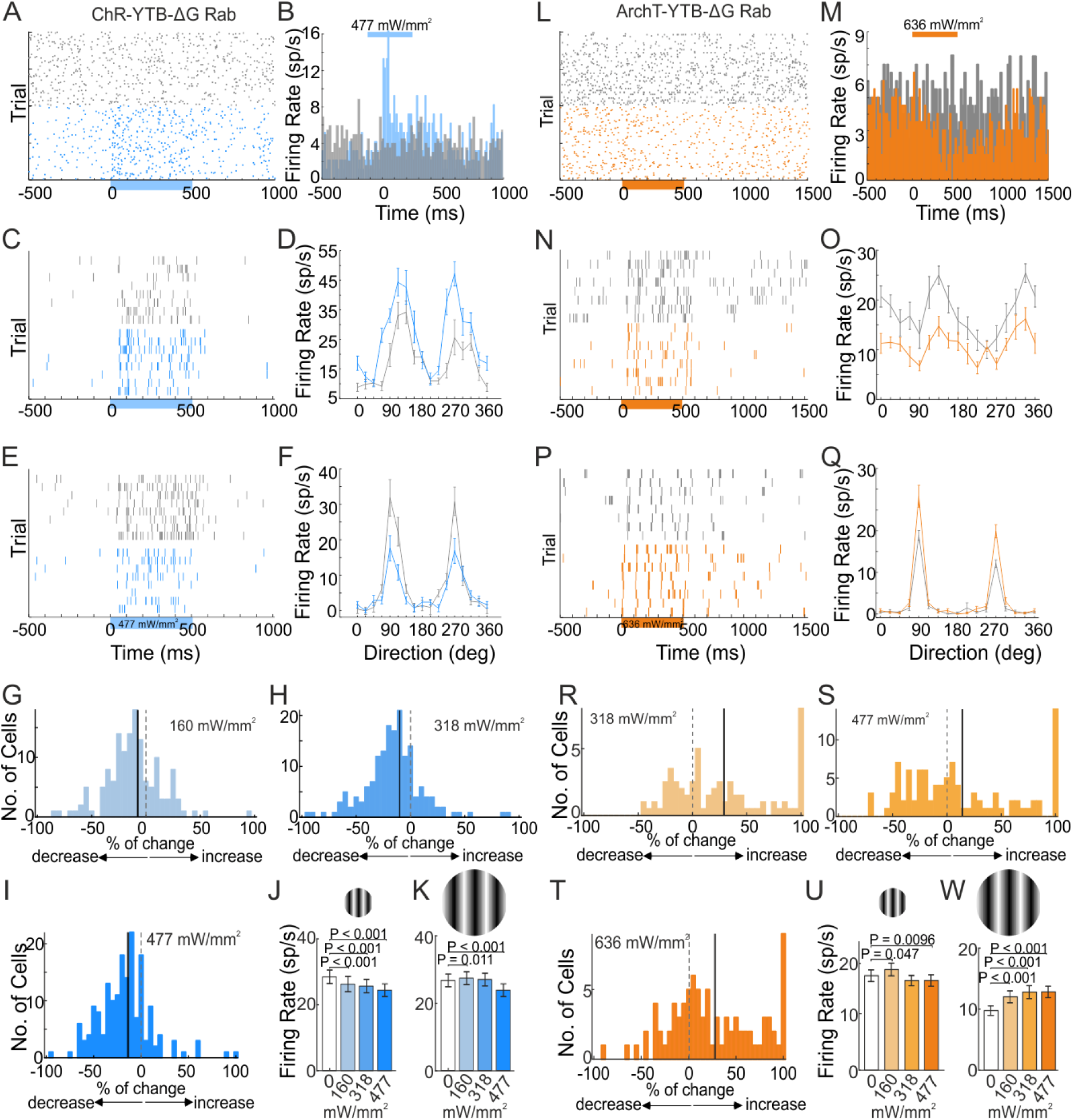
| Bidirectional optogenetic control of firing rate in the PV-related circuit. **a**, Raster of 100 trials of background activity (black) and 100 trials during 477 mW mm⁻² illumination (blue) for a ChR2-expressing neuron, presumed to be a PV interneuron. b, Peristimulus time histogram for the neuron in a; the horizontal line marks the 500-ms epoch of 473-nm illumination. c, Raster showing an increased response to drifting gratings during 477 mW mm⁻² blue illumination. d, Orientation tuning curves for the neuron in c without illumination (black) and with 477 mW mm⁻² blue illumination (blue). e, A neuron representative of the majority of recorded cells, showing a reduced visual response during illumination. f, Tuning curves for the neuron in e. g, Distribution of per-cell percentage change in firing rate at 477 mW mm⁻² relative to baseline; negative values indicate reduced firing. h, Mean firing rate versus illumination power for CRF stimuli. i, As in h, for full-field stimuli. j, Raster of 100 trials of background activity (black) and 100 trials during 636 mW mm⁻² illumination (yellow) for an ArchT-expressing neuron, presumed to be a PV interneuron. k, Peristimulus time histogram for the neuron in j; the horizontal line marks the 500-ms epoch of 589-nm illumination. l, Raster showing a reduced response to drifting gratings during 636 mW mm⁻² yellow illumination in the same presumed ArchT-expressing neuron. m, Tuning curves for the neuron in l without illumination (black) and with 636 mW mm⁻² yellow illumination (orange). n, A neuron representative of the majority of recorded cells, showing an enhanced visual response during illumination. o, Tuning curves for the neuron in n. p, Distribution of per-cell percentage change in firing rate at 636 mW mm⁻² relative to baseline. q, Mean firing rate versus illumination power for CRF stimuli. r, As in q, for full-field stimuli. Data are mean ± s.e.m. P values are from two-tailed Wilcoxon signed-rank tests comparing each illumination power with the no-illumination baseline. Exact n and P values are given in Supplementary Tables 1 and 2. See also Supplementary Fig. 1.

Across the population, ChR2 activation reduced firing rate for CRF stimuli at every illumination power (Fig. 4h; 28.39 ± 28.06 spikes s^−1^ at the pooled baseline, n = 187 neurons; on paired comparisons firing declined to 26.27 ± 28.13 spikes s^−1^, n = 145, P = 2.2 × 10^−6^; 25.65 ± 26.39 spikes s^−1^, n = 171, P = 1.5 × 10^−12^; and 24.41 ± 25.28 spikes s^−1^, n = 177, P = 2.5 × 10^−14^, at 160, 318 and 477 mW mm^−2^, respectively; baseline-subtracted net firing rate showed the same pattern, declining from 22.64 ± 17.21 to 18.69 ± 15.15 spikes s^−1^, P < 0.0001 at every power; Supplementary Table 1). For FF stimuli, firing rate decreased significantly at the lowest and highest powers and was unchanged at the intermediate power (Fig. 4i; 27.03 ± 27.04 spikes s^−1^ at baseline, n = 205; 27.71 ± 27.20 spikes s^−1^, n = 202, P = 4.8 × 10^−3^, at 160 mW mm^−2^; 27.48 ± 27.10 spikes s^−1^, n = 198, P = 0.72, at 318 mW mm^−2^; and 24.10 ± 26.03 spikes s^−1^, n = 188, P = 6.3 × 10^−5^, at 477 mW mm^−2^; Supplementary Table 2).

ArchT activation produced only modest changes in CRF firing rate (Fig. 4q; 17.70 ± 9.91 spikes s⁻¹ at baseline, n = 82, versus 18.86 ± 10.77 spikes s⁻¹, P = 0.048, at 160 mW mm⁻²; 16.79 ± 8.66 spikes s⁻¹, P = 0.073, at 318 mW mm⁻²; 17.25 ± 10.90 spikes s⁻¹, P = 0.15, at 477 mW mm⁻²; and 16.40 ± 9.92 spikes s⁻¹, P = 4.1 × 10⁻³, Holm-corrected P = 0.016, at 636 mW mm⁻²; Supplementary Table 1). For FF stimuli, by contrast, ArchT activation increased firing rate at every power (12.03 ± 8.74 spikes s⁻¹, P = 3.3 × 10⁻⁶, at 160 mW mm⁻²; 12.80 ± 9.58 spikes s⁻¹, P = 2.5 × 10⁻⁷, at 318 mW mm⁻²; 11.17 ± 7.66 spikes s⁻¹, P = 0.034, at 477 mW mm⁻²; and 11.25 ± 8.53 spikes s⁻¹, P = 4.9 × 10⁻⁴, at 636 mW mm⁻²; baseline 9.78 ± 7.08 spikes s⁻¹, n = 82; Fig. 4r and Supplementary Table 2), consistent with release of a suppression that is present only when the surround is stimulated.

When ChR2 was expressed in PV cells alone, rather than in the PV-related circuit, firing-rate effects were weaker and restricted to CRF stimuli (Supplementary Fig. 1j, k; 12.91 ± 8.59 spikes s^−1^ at baseline, n = 54, versus 13.92 ± 9.69 spikes s^−1^, n = 54, P = 0.40, and 14.88 ± 10.61 spikes s^−1^, n = 54, P = 0.040, at 160 and 318 mW mm^−2^, and 12.30 ± 8.43 spikes s^−1^, n = 54, P = 0.32, at 477 mW mm^−2^; net firing rate reached significance at the intermediate power, 13.31 ± 9.95 spikes s^−1^, P = 8.2 × 10^−3^; Supplementary Table 1). FF stimulation produced no sustained population effect in this condition (9.83 ± 11.87 spikes s^−1^ at baseline versus 9.63 ± 13.06 spikes s^−1^ at 477 mW mm^−2^, n = 128, P = 0.20; Supplementary Table 2). Modulation magnitude scaled with illumination power density in all conditions, as expected for direct optogenetic control.

### PV-circuit activation sharpens orientation tuning for classical-receptive-field stimuli

We next measured orientation tuning under the same manipulations. Responses to drifting gratings presented at 16 directions in 22.5° steps were recorded at four illumination powers for ChR2 and five for ArchT (0, 160, 318 and 477 mW mm^−2^ for ChR2; 0, 160, 318, 477 and 636 mW mm−2 for ArchT), fitted with a double Gaussian, and summarized by the half-width at half-height (HWHH) of the Gaussian centred on the preferred direction. Representative CRF tuning curves for ChR2 activation are shown in Fig. 5a and Supplementary Fig. 2a, k, l, o, p, and the corresponding FF curves in Fig. 5f and Supplementary Fig. 2f; equivalent examples for ArchT are shown in Fig. 5k, p.

**Fig. 5.**
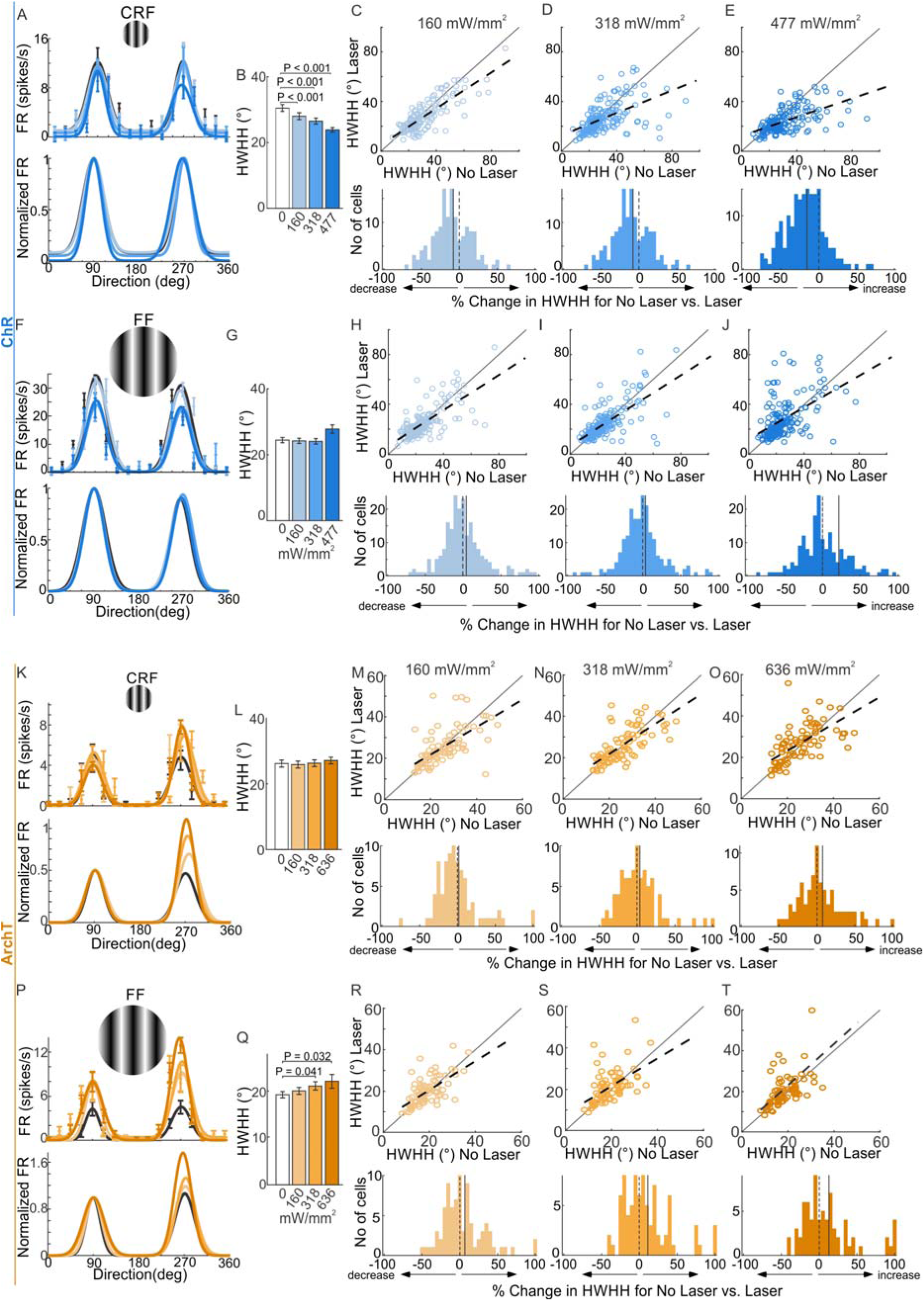
| Optogenetic control of orientation tuning width in the PV-related circuit. Half-width at half-height (HWHH) for the PV-related circuit infected with AAV-PV-YTB and either EnvA-mCherry-ChR2-dG rabies virus (blue) or EnvA-GFP-ArchT-dG rabies virus (yellow), for classical receptive field (CRF) and full-field (FF) stimuli. a, Tuning curves from one neuron at each illumination power at the preferred stimulus size; top, mean responses with s.e.m. and Gaussian fits colour-coded by power; bottom, the same fits normalized. b, Mean CRF HWHH at each illumination power during ChR2 activation. c–e, Per-cell HWHH for no illumination versus 160 (c), 318 (d) and 477 mW mm⁻² (e); top, paired scatter plots; bottom, distributions of percentage change. f, As in a, for full-field stimuli during ChR2 activation. g, Mean FF HWHH at each illumination power during ChR2 activation. h–j, As in c–e, for full-field stimuli. k, As in a, during ArchT activation. l, Mean CRF HWHH at each illumination power during ArchT activation. m–o, Per-cell HWHH for no illumination versus 318 (m), 477 (n) and 636 mW mm⁻² (o). p, As in a, for full-field stimuli during ArchT activation. q, Mean FF HWHH at each illumination power during ArchT activation. r–t, As in m–o, for full-field stimuli; solid lines indicate the mean percentage change. Data are mean ± s.e.m. P values are from two-tailed Wilcoxon signed-rank tests comparing each illumination power with the no-illumination baseline. Exact n and P values are given in Supplementary Tables 1 and 2. See also Supplementary Fig. 2.

Activating the ChR2-expressing PV-related circuit sharpened orientation tuning progressively with illumination power, but only for CRF stimuli (Fig. 5b). On paired comparisons, HWHH declined from 31.41 ± 13.96° to 27.99 ± 12.49° at 160 mW mm⁻² (n = 145, −7.9%, P = 1.0 × 10⁻⁵), from 30.73 ± 13.84° to 26.50 ± 11.86° at 318 mW mm⁻² (n = 171, −8.5%, P = 6.4 × 10⁻⁶), and from 30.54 ± 13.86° to 23.86 ± 8.94° at 477 mW mm⁻² (n = 177, −15.7%, P = 7.7 × 10⁻¹⁵); all three comparisons survive Holm correction (Supplementary Table 1). Suppressing the same circuit with ArchT, in contrast, left CRF tuning width unchanged at every power tested (26.16 ± 9.54° at baseline, n = 82; 25.85 ± 9.59°, P = 0.19, at 160 mW mm⁻²; 26.26 ± 9.24°, P = 0.96, at 318 mW mm⁻²; 26.96 ± 10.80°, P = 0.82, at 477 mW mm⁻²; and 27.11 ± 9.53°, n = 80, P = 0.12, at 636 mW mm⁻²; Fig. 5l and Supplementary Table 1). This asymmetry is what a surround-suppression account predicts: a CRF-sized stimulus does not engage the surround, so there is little PV-mediated suppression available to remove and suppressing the circuit leaves tuning width unaffected, whereas driving the circuit adds inhibition and sharpens tuning. Population distributions of per-cell HWHH change are shown in Fig. 5c–e for ChR2 and Fig. 5m–o for ArchT, and the effects are summarized schematically in Fig. 6.

**Fig. 6.**
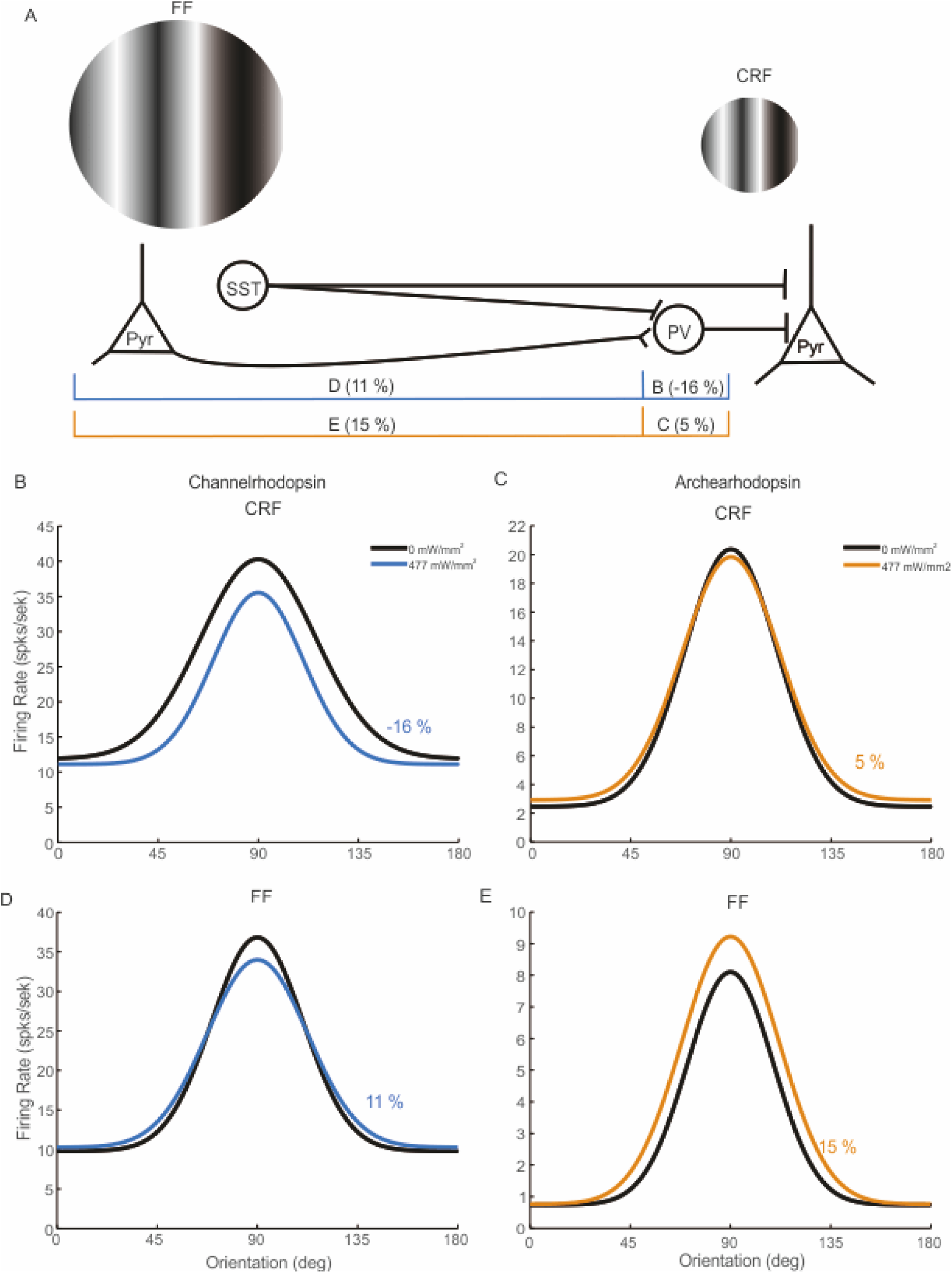
| Summary of the effects of PV-circuit activation and suppression on orientation tuning. **a**, Circuit diagram of the experimental logic and the position of the extraclassical suppressive surround; bracketed values give the mean per-cell change in half-width at half-height at 477 mW mm⁻² for each manipulation and stimulus condition. b, Population-average orientation tuning curve without illumination (black) and during ChR2-mediated activation of the PV-related circuit (blue) for classical-receptive-field stimuli; tuning narrows by 16% (P = 7.7 × 10⁻¹⁵). c, As in b, during ArchT-mediated suppression (orange) for classical-receptive-field stimuli; tuning width is unchanged (+5%, P = 0.82). d, As in b, for full-field stimuli; the change is not significant (+11%, P = 0.095). e, As in c, for full-field stimuli; tuning broadens by 15% (P = 0.0078, Holm-corrected P = 0.031). Curves are Gaussians constructed from the population mean half-width and peak rate for each condition. Activation therefore sharpens tuning only where the surround is not engaged, and suppression broadens it only where the surround is engaged. Asterisks mark comparisons significant after Holm correction; n.s., not significant.

As a fit-free corroboration, we also computed the orientation-selectivity index (OSI), a global vector measure that does not depend on the Gaussian fit. In the PV-related ChR2 cohort, OSI increased with illumination power for CRF stimuli, mirroring the HWHH sharpening: +13.5%, +18.5% and +24.1% at 160, 318 and 477 mW mm^−2^ (P = 0.069, 2.3 × 10^−3^ and 1.2 × 10^−5^; the latter two survive Holm correction; Supplementary Table 1). For FF stimuli, OSI did not increase at any power and decreased slightly at the highest power (−4.7%, P = 6.9 × 10^−5^; Supplementary Table 2), consistent with the tuning-width result and independent of the fitting procedure.

### Surround engagement occludes PV-dependent sharpening

Tuning measured with surround-engaging stimuli behaved differently. First, baseline tuning was already substantially narrower with FF than with CRF stimuli in both cohorts (24.1 ± 11.1° versus ∼30–31° in the ChR2 cohort; 19.2 ± 6.1° versus 26.2 ± 9.5° in the ArchT cohort), consistent with previous reports that surround stimulation sharpens orientation tuning in cat V1^21,23^. Second, ChR2 activation did not sharpen this already-narrow tuning at any power. FF HWHH was statistically unchanged across the range tested (24.13 ± 11.14° at baseline, n = 205; 23.91 ± 10.81°, n = 202, P = 0.99, at 160 mW mm^−2^; 23.87 ± 11.14°, n = 198, P = 0.92, at 318 mW mm^−2^; and 27.30 ± 17.53°, n = 188, P = 0.095, at 477 mW mm^−2^; Supplementary Table 2). The apparent broadening of FF tuning at the highest ChR2 power did not reach significance on paired testing and is not interpreted here. Per-cell distributions are shown in Fig. 5h–j and Fig. 5r–t.

For the suppression arm, the pattern reversed. ArchT broadened FF tuning — the condition in which the surround, and with it PV-mediated surround suppression, is engaged. HWHH increased from 19.20 ± 6.09° at baseline (n = 82) to 20.01 ± 6.66° (P = 0.29) at 160 mW mm⁻², 21.05 ± 7.65° (+9.6%, P = 0.040) at 318 mW mm⁻² and 22.03 ± 8.76° (+14.7%, P = 0.0078, Holm-corrected P = 0.031) at 477 mW mm⁻², and remained broadened at 636 mW mm⁻² (22.00 ± 12.14°, n = 72, P = 0.065; Fig. 5q and Supplementary Table 2). Removing PV-mediated inhibition therefore widens orientation tuning specifically when the surround is engaged, the complement of the sharpening that driving the circuit produces when it is not.

The contrast between stimulus conditions is therefore one of occlusion of the activation-dependent sharpening. Within the classical receptive field, tuning width narrows with PV-circuit activation but is unchanged by suppression. Once the surround is engaged, the relationship reverses: activation no longer sharpens tuning, whereas suppression broadens it. This is the signature expected if surround suppression and PV-mediated inhibition converge on a common inhibitory mechanism that is already close to saturation when the surround is stimulated, so that the sharpening available within the classical receptive field is occluded once surround circuitry sets tuning width. The broadening of FF tuning under ArchT suppression is considered in the Discussion.

### Restricting expression to PV cells alone produces weaker sharpening

To assess how much of the effect required the presynaptic partners of PV cells rather than PV cells themselves, we repeated the activation experiment with ChR2 expression restricted to PV interneurons. Sharpening of CRF tuning was present but weaker and reached significance only at the highest power: HWHH declined from 34.15 ± 12.44° (n = 50) to 31.22 ± 10.84° at 160 mW mm^−2^ (−4.2%, P = 0.079), 31.24 ± 11.26° at 318 mW mm^−2^ (n = 47, −4.6%, P = 0.035), and 29.54 ± 11.88° at 477 mW mm^−2^ (n = 49, −9.2%, P = 9.1 × 10^−4^, Holm-corrected P = 0.006). This is the same power-dependent sharpening seen in the full PV-related circuit, but with roughly half the magnitude, indicating that presynaptic partners of PV cells are a major contributor to the effect. FF tuning was unaffected at every power in this condition (30.14 ± 18.85° at baseline versus 28.69 ± 18.96°, n = 123, P = 0.30, at 160 mW mm^−2^; 32.83 ± 26.11°, n = 121, P = 0.46, at 318 mW mm^−2^; and 31.64 ± 23.32°, n = 124, P = 0.33, at 477 mW mm^−2^; Supplementary Table 2 and Supplementary Fig. 2). Mean changes in HWHH at the highest illumination power tested are summarized in Fig. 6b–e: −16% for ChR2 with CRF stimuli and +15% for ArchT with full-field stimuli, computed as the mean of per-cell percentage changes; the complementary comparisons were not significant (+5% for ArchT with CRF stimuli, +11% for ChR2 with full-field stimuli; Fig. 6c,d).

Together, these results show that the PV-related circuit controls orientation tuning width bidirectionally within the classical receptive field, that this control is stronger when the presynaptic partners of PV cells are co-recruited than when PV cells are driven alone, and that the activation-dependent sharpening does not extend to the already-sharpened tuning measured when the extraclassical surround is engaged.

## Discussion

PV-mediated sharpening of orientation tuning in cat V1 depended on stimulus size. Within the classical receptive field, activating the PV-related circuit narrowed tuning in proportion to illumination power, whereas suppressing it left tuning unchanged. Once stimuli extended into the extraclassical surround, the pattern reversed: activation no longer sharpened tuning, whereas suppression broadened it. This double dissociation places PV circuitry in the causal path that sets tuning width, and identifies surround suppression as the inhibition it supplies. The limit on activation-dependent PV sharpening is therefore set by the receptive-field architecture through which PV cells are recruited, not by a fixed property of the PV population. This offers a parsimonious account of why optogenetic manipulations of PV neurons have yielded reports of divisive gain control in some studies and sharpening in others: the discrepancy may reflect differences in cortical architecture and stimulus context rather than differences in PV physiology.

PV interneurons receive robust, largely non-selective feedforward excitation from the thalamus, and in layer 4 this geniculocortical drive recruits them rapidly, producing widespread inhibition of nearby excitatory neurons^30^. Because the inhibition closely follows thalamic excitation, it narrows the window over which cortical neurons integrate their inputs^31^, enhancing temporal precision and stimulus selectivity; consistent with this, pharmacological blockade of GABA_A_ receptors broadens orientation tuning in pyramidal neurons^32^. On its own, however, this mechanism predicts sharpening regardless of stimulus size, because thalamic recruitment of PV cells is engaged by stimuli confined to the classical receptive field. Our finding that PV manipulation changes tuning width within the classical receptive field but that activation no longer sharpens once the surround is engaged therefore argues that feedforward recruitment is not the whole explanation.

A second candidate lies in the recurrent connectivity between PV interneurons and local pyramidal neurons. Individual PV cells receive convergent input from excitatory neurons spanning a wide range of orientation preferences and therefore respond broadly^15,33^, encoding overall excitatory drive rather than any particular feature. On this view, raising PV activity should suppress cortical output divisively, scaling firing rates without markedly changing tuning width^1,3,34^, as in the canonical normalisation model^31^. Our firing-rate data are consistent with divisive action, but the accompanying change in tuning width within the classical receptive field indicates that divisive normalisation alone does not capture the effect of PV recruitment in a columnar cortex.

Not all PV neurons conform to this broadly tuned profile; some are as orientation-selective as excitatory cells, reflecting more structured local connectivity^35,36^. This may be where cortical architecture matters most. In columnar species such as cats and primates, an individual PV cell integrates input across several orientation domains, whereas in rodents, which lack orientation maps, local circuit specificity can leave a subset of PV cells comparatively sharply tuned^35^. A surround stimulus in a columnar cortex recruits inhibition across many domains at once, and the broadly acting PV population it engages is precisely the one whose further activation we found to produce no additional sharpening.

The broadening of full-field tuning under ArchT suppression is the complement of the sharpening seen within the classical receptive field, and is what a surround-suppression account predicts: removing PV-mediated inhibition widens tuning precisely in the condition in which that inhibition is recruited. It reached significance at 477 mW mm⁻² (+14.6%, P = 0.0078, Holm-corrected P = 0.031) and remained present at the highest power tested (+12.9%, P = 0.065). Because full-field responses are of lower rate and their half-width estimates correspondingly noisier, we treat the magnitude as approximate and rest the mechanistic claim on the direction of the effect and on its stimulus specificity rather than on its size.

Beyond local circuitry, PV interneurons are gated by top-down feedback and by neuromodulation, either of which could contribute to a surround-specific role. Feedback projections from higher visual areas engage PV cells directly and are well placed to regulate surround suppression and contextual integration^37^. Cholinergic input adds a further layer: in primates, most PV neurons express muscarinic receptors, and acetylcholine enhances their inhibitory output without altering orientation preference^38^, while nicotinic input adjusts network gain in macaque V1^39^. Vasoactive intestinal peptide-expressing interneurons transiently suppress PV activity during locomotion and attention, disinhibiting pyramidal neurons^40^. We did not manipulate feedback or neuromodulatory state, but their capacity to gate PV output in a context-dependent manner is a plausible route by which surround engagement, rather than PV identity alone, could set the limits of sharpening.

PV-circuit activation also has consequences that single-neuron tuning width does not capture. Driving fast-spiking PV cells induces gamma rhythms that improve temporal precision and perceptual discrimination^41,42^, synchronising pyramidal activity and coordinating ensembles engaged in attention and sensory selection. The sharpening we observed may therefore be accompanied by changes in network timing that our recordings were not designed to resolve.

Several limitations bound these conclusions. First, the rabies-virus strategy delivers the actuator to PV interneurons and to the neurons presynaptic to them, which include excitatory cells; illuminating this network is therefore not equivalent to activating PV cells selectively, and the direct comparison with PV-restricted expression, in which sharpening was weaker and not sustained, is the relevant control rather than an incidental one.

Second, the illumination powers used here are high. The opposite-signed effects of ChR2 and ArchT on tuning width-narrowing under activation with classical-receptive-field stimuli, broadening under suppression with surround-engaging stimuli-argue against a thermal or light-artefact explanation, because a passive artefact would be expected to act in the same direction regardless of the opsin and of stimulus size; we did not, however, record from uninfected cortex under matched illumination, and such a control-from a pre-injection session, a contralateral hemisphere, or a site outside the injection core-would settle the point directly. Third, because the tuning model includes an explicit baseline term, additive changes in spontaneous activity are absorbed by that term and divisive scaling of the response leaves the fitted width σ unchanged; the observed changes in HWHH therefore cannot be attributed to changes in response gain or background firing, a conclusion corroborated by the fit-free OSI result. Fourth, cells were not held across all illumination powers, so the numbers of neurons differ between conditions, and comparisons are paired only within cells common to a given pair. Finally, recordings were made under anaesthesia, which attenuates the top-down and neuromodulatory influences discussed above.

Taken together, these results indicate that PV interneurons shape orientation selectivity not through a single fixed action but through a repertoire determined by which circuit a stimulus engages: feedforward inhibition limits the integration window, recurrent connectivity supports divisive normalisation, and top-down and neuromodulatory pathways adjust inhibitory tone with behavioural state. The presence of orientation columns in carnivores and primates may determine which of these dominates for a given stimulus configuration, favouring surround-dependent sharpening over uniform gain control. Because disrupting PV function degrades response precision while enhancing it can improve visual discrimination^34^, the mechanism matters functionally: the PV circuit acts less as a static gain controller than as a context-dependent regulator of selectivity whose net effect on tuning is set by the receptive-field architecture, and in particular the extraclassical surround, through which it is recruited.

## Methods

### Animals and ethical approval

Experiments were performed in vivo on seven adult domestic cats (*Felis catus*), aged 8–24 months and weighing 2.4–5 kg, of either sex, obtained from an institutionally approved colony. All procedures complied with the guidelines of, and were approved by, the Institutional Animal Care and Use Committee and the Institutional Biosafety Committee of the University of California, Irvine, followed National Institutes of Health guidelines, and are reported in accordance with the ARRIVE 2.0 guidelines. The seven animals were distributed across the three viral conditions (ChR2 PV-related circuit, ArchT PV-related circuit and ChR2 PV-only).

### Generation of AAV-PV-YTB

Recombinant AAV particles encoding YTB under the control of a PV promoter (Fig. 1a) were produced by removing the 3.1-kb human GAD1 promoter fragment from AAV-GAD1-YTB^28^ and replacing it with the 2.9-kb pufferfish PV promoter^29^. The PV promoter was amplified with the primers 5[-ggcggacgcgtcgaattcgattttgaccgtttg-3[and 5[-ggcggggcgcgcctttgccagctgaaaagacagg-3[using PCSC-SP-PW-GFP FuPV as template (primers synthesized by Integrated DNA Technologies). Each reaction contained 45 µl SuperMix (Thermo Fisher Scientific), 2 µl of each primer at 10 pmol µl^−1^ and 1 µl template DNA at 10 ng µl^−1^, and was cycled at 95 °C for 10 min followed by 32 cycles of 95 °C for 30 s, 52 °C for 30 s and 72 °C for 3 min. The purified product was digested with AscI and MluI to release the PV promoter; AAV-GAD1-YTB^28^ was digested with the same enzymes to remove the GAD1 promoter and gel-purified. Vector and insert were ligated and transformed into chemically competent *Escherichia coli* (Thermo Fisher Scientific). To increase expression, the YTB transgene is preceded by a 525-bp intronic splice donor and acceptor sequence from human beta-globin and followed by a 264-bp bovine growth hormone polyadenylation signal (Fig. 1a). The resulting plasmid, AAV-PV-YTB, was packaged as AAV9 by the Salk Institute viral vector core at a titre of 1.33 × 10^11^ genome copies ml^−1^. Following our previous work^28^, the AAV was diluted to 25% of stock titre to optimize cell-type specificity.

### Viral injections

For injection of the AAV-PV-YTB or AAV-PV-YTx helper virus, cats were anaesthetized with ketamine (20 mg kg^−1^, i.m.) and xylazine (3 mg kg^−1^, i.m.) and maintained on isoflurane, then placed in a stereotaxic frame. A craniotomy was made under sterile conditions over the area 17/18 border, near the midline of the caudal third of the dorsolateral cortical surface, and the dura was removed. Virus was pressure-injected through a glass pipette with a tip broken to approximately 20 µm, lowered with a computer-controlled micropositioner on a stereotaxic arm. Injections of 0.3–1.5 µl were made at several V1 locations at three cortical depths (600, 1,200 and 1,800 µm). For animals scheduled to receive rabies virus, the anteroposterior and mediolateral coordinates of each vector-injected site were recorded and pencil landmarks were placed at four positions around the edge of the craniotomy to allow the site to be relocated approximately three weeks later. The craniotomy was then covered with artificial dura (Tecoflex, Microspec), sealed with dental acrylic, and the animal was recovered. Approximately three weeks later, EnvA-dG-RV-mCherry-ChR2 or EnvA-dG-RV-GFP-ArchT (0.4 µl) was injected as close as possible to the original site, guided by the recorded coordinates, the pencil landmarks and the surface vasculature. The craniotomy was resealed and the animal recovered for a further 7–14 days before acute electrophysiology.

### Animal preparation for electrophysiology

Animals were anaesthetized with ketamine (21 mg kg^−1^, i.m.) and xylazine (3 mg kg^−1^, i.m.), after which tracheal and venous cannulation were performed. Anaesthesia was maintained with isoflurane (0.2–1.0%) in a 2:1 mixture of nitrous oxide and oxygen. Depth of anaesthesia was monitored throughout using lung pressure, electrocardiogram and electroencephalogram. End-tidal CO_2_ was held between 4.0 and 4.5% and body temperature near 37 °C with a homeothermic blanket (Harvard Apparatus). Pupils were dilated and accommodation blocked with 1% atropine sulfate, and the corneas were protected with air-permeable corrective contact lenses. A craniotomy was made over the dorsal surface of area 17 and the dura removed. Eye movements were prevented by neuromuscular blockade with gallamine triethiodide (40 mg in 1 ml, i.v.), maintained at 2.0 mg kg^−1^ h^−1^ in 5% dextrose and lactated Ringer’s solution containing dexamethasone (0.5 mg kg^−1^ h^−1^, i.v.).

### Electrophysiology and visual stimulation

Single-unit recordings were made with epoxy-insulated tungsten microelectrodes (3–7 MΩ; FHC) and 24-channel linear U-probes (Plexon). Electrodes were advanced perpendicular to the cortical surface 3–4 mm from the midline, and recording depth was used to assign units to supragranular (<600 µm), granular (600–1,200 µm) and infragranular (>1,200 µm) compartments as described previously^20^. After insertion, the cortex was covered with 1.5% agar in saline and sealed with physiological wax to reduce pulsation. Signals were acquired with a 32-channel Scout system (Ripple), band-pass filtered from 500 Hz to 7 kHz and digitized at 30 kHz. Spikes were sorted online in Trellis (Ripple) and offline with Plexon Offline Sorter. Visual stimuli were generated in MATLAB (MathWorks) with Psychophysics Toolbox 3^43–45^ and displayed on a gamma-calibrated 21-inch ViewSonic G225f CRT monitor at a mean luminance of 50 cd m^−2^, 640 × 480 pixels and 100 Hz. Stimulus onset times were corrected for monitor delay using a photoresistor and an Arduino Zero microcontroller. Each cell was stimulated through the dominant eye with the non-dominant eye occluded.

Once a unit was isolated, the approximate receptive-field centre and preferred orientation were found using a circular square-wave grating patch approximately 4° in diameter; the patch was then reduced to 1° or less and systematically moved to locate the centre, as described previously^17,20,46^. Receptive-field eccentricities ranged from 2° into the upper visual quadrant to 16° into the lower quadrant. Preferred orientation was then reassessed with drifting sine-wave gratings sampled at 16 directions in 22.5° steps within a circular aperture approximately 10° in diameter at 100% contrast, 0.2 cycles deg^−1^ and 4 Hz, after which optimal spatial and temporal frequency were determined. Using all optimal parameters at 100% contrast, responses were measured across 18 aperture sizes from 0.2° to 30°. From this size-tuning curve, the CRF was defined as the aperture producing the largest mean response and the extraclassical surround as the larger aperture producing the greatest subsequent response reduction. Orientation tuning was then measured at optimal spatial and temporal frequency with CRF-sized and surround-engaging (full-field) stimuli at high contrast. All stimuli were drifting sinusoidal gratings in sharp-edged circular apertures on a grey background of matched mean luminance, presented for 0.5–0.8 s and repeated eight times with an inter-stimulus interval of 1.5 s; a 100-ms blank preceding each presentation was used to estimate spontaneous activity.

### Optogenetic stimulation

A 200-µm-diameter optical fibre (0.39 numerical aperture) was inserted alongside the recording electrode. Illumination was controlled and synchronized with visual stimulation by a Master-8 stimulator (AMPI). Optogenetic manipulation began after optimal visual parameters had been identified, and a separate recording was made at each illumination power, with 2-min intervals between conditions to allow baseline activity to recover. Power densities at the fibre tip were 0, 160, 318 and 477 mW mm^−2^ for ChR2 experiments (473 nm) and 0, 160, 318, 477 and 636 mW mm−2 for ArchT experiments (589 nm). The illumination epoch spanned the visual stimulus and was 500 ms, synchronized to visual-stimulus onset.

### Data analysis

Tuning curves were computed from mean spike rates. The orientation-selectivity index (OSI) was calculated as

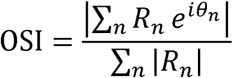

where is the θ*_n_* is the *n*th stimulus orientation and the corresponding response. Orientation responses were fitted with Gaussian functions^16,47,48^ of the form

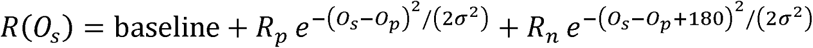

Where *O_s_* is stimulus orientation, R(*O_s_*) the response, the preferred orientation *R_p_* and *R_n_* the responses at the preferred and non-preferred orientations, σ the tuning width and ‘baseline’ the offset. Fits were made without subtracting spontaneous activity, following Alitto and Usrey^48^ and our previous work^49^. Orientation tuning bandwidth was taken as the half-width at half-height (HWHH), with HWHH = 1.18 × σ. Fits were accepted when R^2^ exceeded 0.9.

### Statistics and reproducibility

Population comparisons used the two-tailed Wilcoxon signed-rank test for paired samples, comparing each illumination power with the no-illumination baseline recorded in the same session. Because several illumination powers were compared with a common baseline within each cohort and stimulus condition, P values were additionally corrected for multiple comparisons using the Holm–Bonferroni procedure within each cohort and measure family; both uncorrected and corrected P values are reported in Supplementary Tables 1 and 2. P values quoted in the text are uncorrected unless stated; the comparisons that survive Holm correction are the three ChR2 CRF tuning-width comparisons, the PV-only CRF comparison at 477 mW mm−2, and the ArchT full-field tuning-width comparison at 477 mW mm−2, together with the firing-rate and orientation-selectivity effects indicated in Supplementary Tables 1 and 2; correction does not remove any of the tuning-width results on which the conclusions rest. Baselines quoted in the text are the paired-subset baselines for each comparison rather than a pooled baseline, so that each mean ± s.d. refers to the same cells as the laser condition with which it is compared. Values in the text are mean ± s.d. and error bars in figures are s.e.m., as indicated in the legends. Exact numbers of neurons (paired counts) and exact P values for every comparison are given in Supplementary Tables 1 (CRF) and 2 (FF). Neurons that were not held for the full sequence of illumination powers contribute only to the comparisons for which both members of the pair were recorded, which is why n differs between powers.

Recordings were obtained from seven animals. To guard against pseudoreplication arising from treating neurons as independent, the number of neurons contributed by each animal in each condition is reported in Supplementary Table 1, and the primary CRF tuning-width comparisons were additionally examined with a linear mixed-effects model with illumination power as a fixed effect and animal as a random intercept; the fixed-effect estimates are reported alongside the Wilcoxon results in Supplementary Table 1. No statistical method was used to predetermine sample size; the number of recorded neurons was determined by the yield of the acute preparations. Experiments were not randomized and investigators were not blinded to viral condition, since the condition is determined by the injected construct. All analyses were performed in MATLAB (MathWorks).

### Histology

At the end of the terminal recording session animals were deeply anaesthetized with sodium pentobarbital (Euthasol) and perfused transcardially with saline followed by 4% paraformaldehyde in 0.1 M phosphate buffer (PB, pH 7.4), as described for the parent cell-type-specific tracing method^28^. Brains were removed and cryoprotected in 30% sucrose in0.1 M PB for approximately 48 h before sectioning.

Cat hemispheres were sectioned coronally at 30 µm for up to 1 mm anterior and posterior to the V1 injection site, to improve PV antibody penetration, and at 40 µm elsewhere. The imaged V1 region was separated from the remainder of the hemisphere and sectioned tangentially at 30 µm after an initial 120-µm section that preserved the surface vascular pattern. Three series of every fourth section were processed for anti-PV immunofluorescence (rabbit polyclonal PV-28, Swant, RRID:AB_2315235) or for DAPI. Co-localization of PV immunofluorescence with YFP was used to establish the cell-type specificity of the AAV helper virus and to identify YFP/mCherry double-labelled starter cells and mCherry-only presynaptic neurons, using a Zeiss Axioplan microscope with 10× (0.45 NA) and 20× (0.8 NA) objectives, with reconstruction in Neurolucida (MBF Bioscience). To limit bleaching, images were captured with a cooled monochrome camera (Cooke SensiCam QE) under Neurolucida control. Two of the four sections in each series were analysed. Free-floating sections were processed on a shaker (100 r.p.m.) at room temperature unless otherwise stated. Sections were rinsed in 1× PBS for 10 min, permeabilized in 0.3% Triton X-100 (Sigma, cat. no. T9284) in PBS for 60 min and blocked for 60 min in 10% normal goat serum (Gibco, cat. no. 16210064), 1% bovine serum albumin (Sigma, cat. no. A7906) and 0.1% Triton X-100 in PBS. Sections were then incubated overnight at 4 °C with the anti-parvalbumin antibody diluted 1:1,000 in PGB diluent (2% normal goat serum, 1% bovine serum albumin and 0.02% Triton X-100 in PBS). After three 10-min rinses in 0.05% Triton X-100 in PBS, immunopositive neurons were revealed with Alexa Fluor 350 goat anti-rabbit IgG (1:500; Invitrogen, cat. no. A-11046) for 2 h at room temperature in the dark, as described previously^28^. Sections were rinsed three times for 10 min in 0.05% Triton X-100 in PBS, washed once in 1× PBS for 10 min, mounted on Superfrost Plus slides (Fisher, cat. no. 12-550-15) and coverslipped with Vectashield mounting medium containing DAPI (Vector Laboratories, cat. no. H-1200) or with Aqua-Mount (Lerner Laboratories, cat. no. 13800).

### Reporting summary

Further information on research design is available in the Nature Portfolio Reporting Summary linked to this article.

### Data availability

The data supporting the findings of this study-per-neuron orientation-tuning bandwidth (HWHH), orientation-selectivity index, firing rate and net firing rate for each viral condition, stimulus size and illumination power-will be deposited on acceptance in the Repository for Open Data (RepOD; https://repod.icm.edu.pl) under DOI [to be assigned on acceptance]. Source data for Figs. 4–6 and Supplementary Figs. 1 and 2 are provided with this paper.

### Code availability

Custom MATLAB code used for orientation-tuning-curve fitting, half-width-at-half-height and orientation-selectivity-index extraction, and the Python code used to regenerate Supplementary Tables 1 and 2 from the per-cell source files and to compute the paired Wilcoxon and Holm-corrected statistics, will be made available in a public GitHub repository on acceptance, with an archived snapshot deposited in the Repository for Open Data (RepOD; https://repod.icm.edu.pl) under DOI [to be assigned on acceptance].

## Acknowledgements

This work was supported by the National Eye Institute (R01EY024890 to D.C.L.); the International Centre for Translational Eye Research (FENG.02.01-IP.05-T005/23), carried out within the International Research Agendas programme of the Foundation for Polish Science and co-financed by the European Union under the European Funds for Smart Economy 2021–2027; the European Union through the project ‘Translational Research and Innovation in Ophthalmology Vision – Centre of Excellence’ (TRIO-VI CoE, no. 101136570), funded under the European Commission’s Teaming for Excellence programme; and the National Science Centre, Poland (2019/34/E/NZ5/00434 to A.T.F.). We thank the Salk Institute viral vector core for virus production. During preparation of this manuscript, the authors used a generative AI tool (Claude, Anthropic) solely to improve the clarity and structure of the written text and to assist with reconciliation and tabulation of the summary statistics from the authors’ own per-cell source data; no AI tool was used to generate, analyse or interpret primary data, or to produce any figure. The authors reviewed and edited all AI-assisted output and take full responsibility for the article’s content.

## Author contributions

A.T.F. and D.C.L. conceived the study. A.T.F., Y.L. and D.C.L. designed the experiments. A.T.F., Y.L., H.S., G.L., L.Z., L.R.S. and R.J. performed the experiments. A.T.F. and Y.L. analysed the data. A.T.F. and D.C.L. wrote the manuscript with input from all authors. D.C.L. and A.T.F. obtained funding. D.C.L. supervised the study.

## Supplementary information

**Supplementary Fig. 1.**
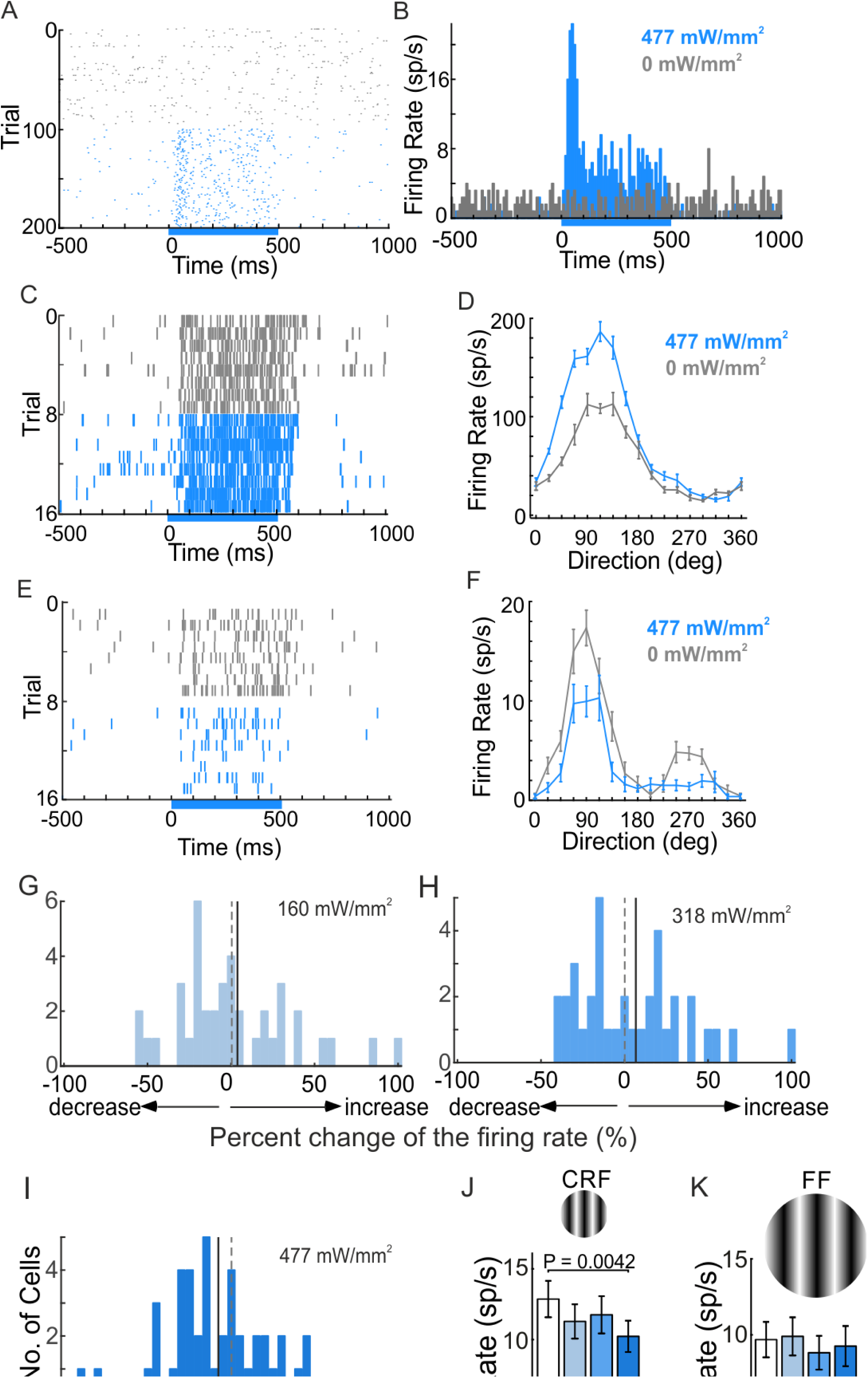
| Optogenetic activation of the PV-only circuit in vivo. **a**, Raster of 100 trials of background activity (black) and 100 trials during 477 mW mm⁻² illumination (blue) for a ChR2-expressing neuron, presumed to be a PV interneuron. b, Peristimulus time histograms before (black) and during (blue) 477 mW mm⁻² illumination. c, Raster showing an increased response to drifting gratings during 477 mW mm⁻² blue illumination. d, Tuning curves for the neuron in c without illumination (black) and with illumination (blue). e, A neuron representative of the majority of recorded cells, showing a reduced visual response during illumination. f, Tuning curves for the neuron in e. g–i, Distributions of per-cell percentage change in firing rate at 160 (g), 318 (h) and 477 mW mm⁻² (i) for the PV-only circuit infected with the ChR2-YTx virus; solid lines mark the distribution mean and dashed lines zero. j, Population mean ± s.e.m. firing rate for CRF stimuli. k, Population mean ± s.e.m. firing rate for full-field stimuli. P values are from two-tailed Wilcoxon signed-rank tests comparing each illumination power with the no-illumination baseline.

**Supplementary Fig. 2.**
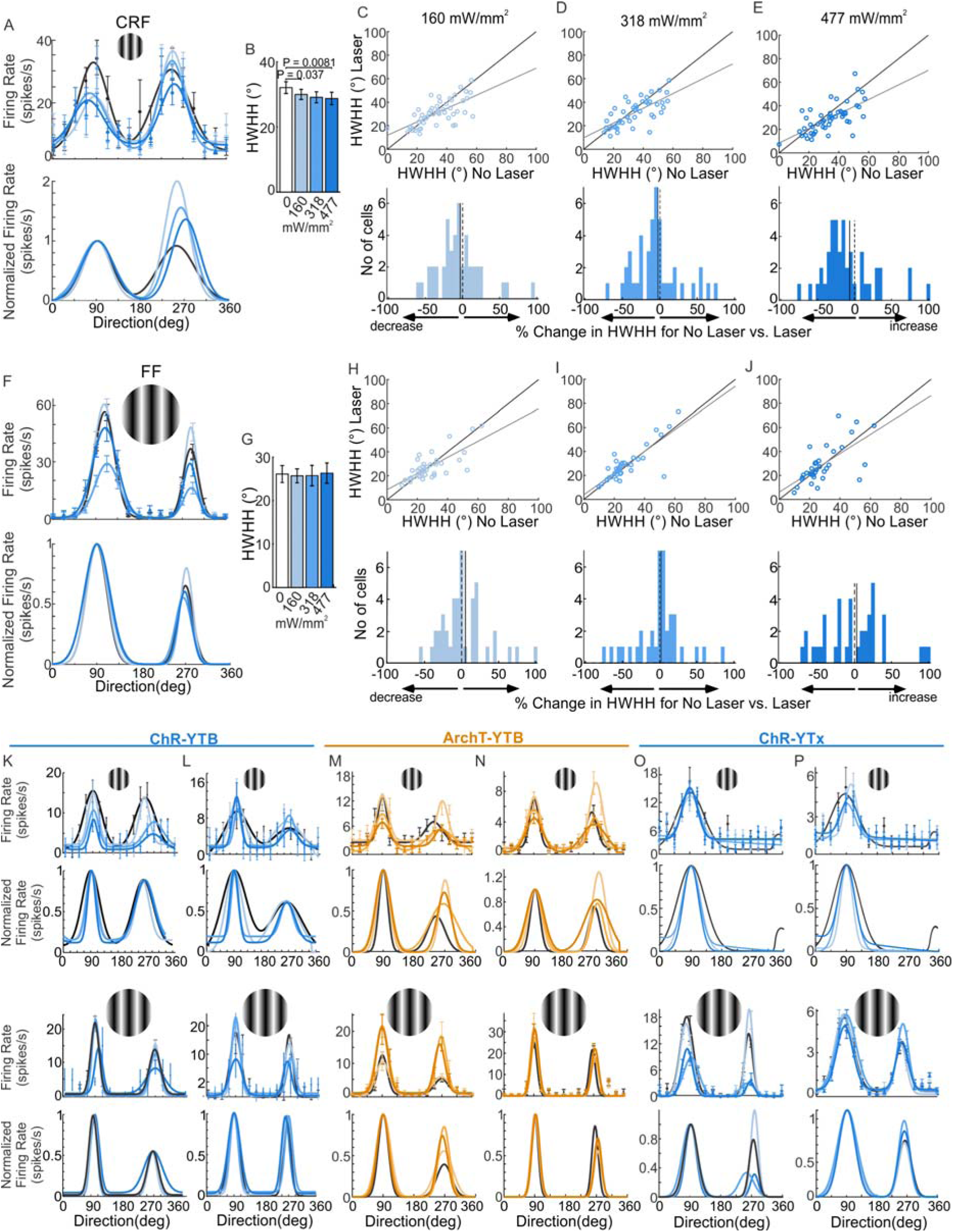
| Orientation tuning width in the PV-only circuit. Half-width at half-height (HWHH) for the circuit infected with AAV-PV-YTx and EnvA-mCherry-ChR2-dG rabies virus, for classical receptive field (CRF) and full-field (FF) stimuli. a, Tuning curves from one neuron at each illumination power at the preferred stimulus size; top, mean responses with s.e.m. and Gaussian fits colour-coded by power; bottom, the same fits normalized. b, Mean CRF HWHH at each illumination power during ChR2 activation. c–e, Top, per-cell CRF HWHH at each of three illumination powers versus no illumination; bottom, distributions of percentage change from the no-illumination condition at 160, 318 and 477 mW mm⁻². Negative values indicate sharpening and positive values broadening. f, As in a, for full-field stimuli. g, Mean FF HWHH at each illumination power during ChR2 activation. h–j, As in c–e, for full-field stimuli; the grey solid line marks the distribution mean and the dashed line marks zero. k–p, Additional single-neuron examples. k,l, Top two rows, mean orientation tuning curves and fits for two neurons recorded at the optimal stimulus size without and with blue illumination (160, 318 and 477 mW mm⁻²) after AAV-PV-YTB and EnvA-mCherry-ChR2-dG injection, with normalized fits beneath; bottom two rows, the same for surround-engaging stimuli. m,n, As in k,l, after AAV-PV-YTB and EnvA-GFP-ArchT-dG injection. o,p, As in k,l, after AAV-PV-YTx and EnvA-mCherry-ChR2-dG injection. Data are mean ± s.e.m. P values are from two-tailed Wilcoxon signed-rank tests comparing each illumination power with the no-illumination baseline.

**Supplementary Fig. 3.**
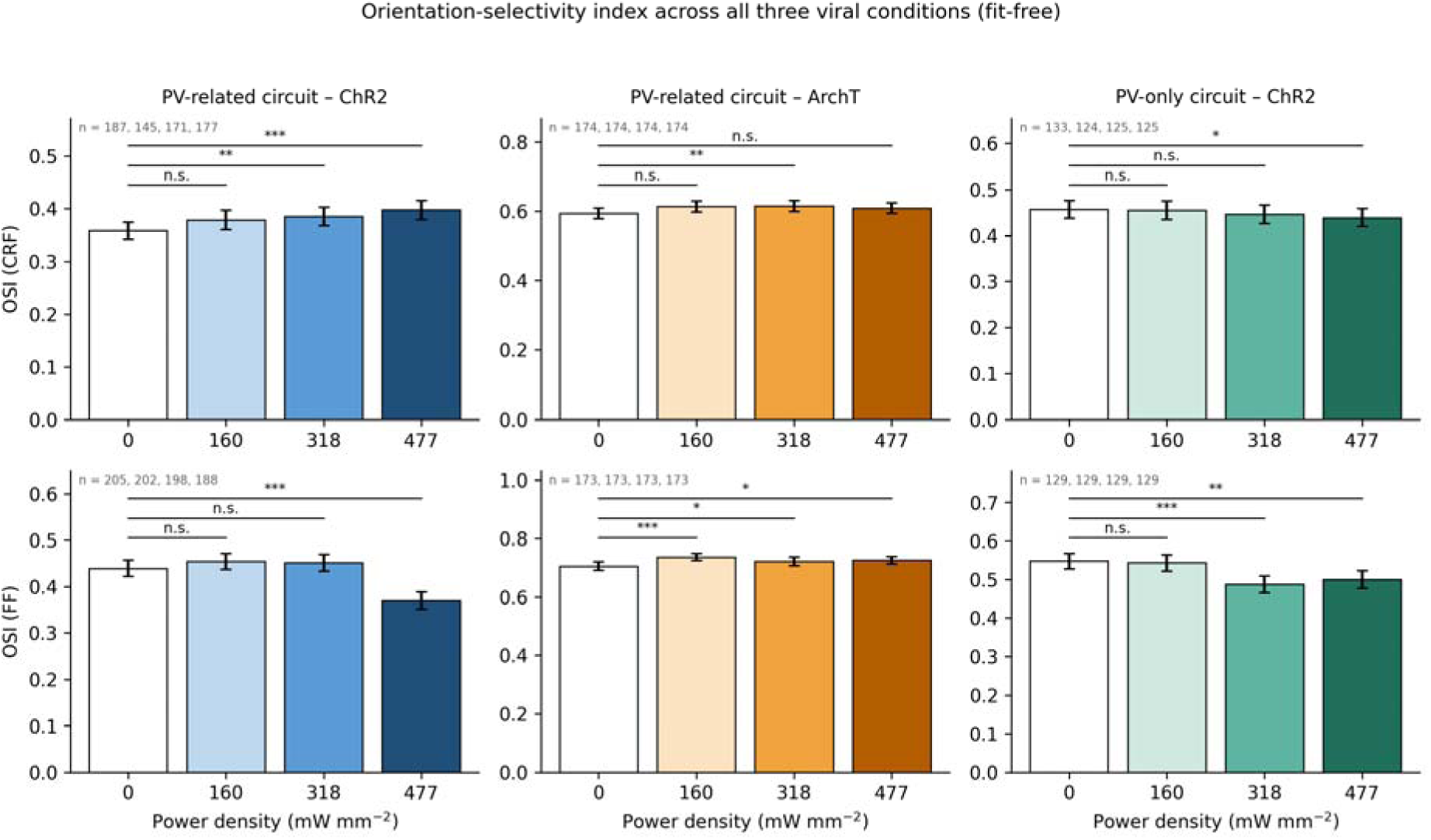
| Orientation-selectivity index across all three viral conditions. Mean orientation-selectivity index (OSI) ± s.e.m. against illumination power density for classical-receptive-field (CRF; top row) and full-field (FF; bottom row) stimuli, for the PV-related circuit expressing ChR2 (left), the PV-related circuit expressing ArchT (middle) and the PV-only circuit expressing ChR2 (right). The OSI is computed directly from the measured responses and does not depend on the Gaussian fit, and so provides a fit-free test of the tuning-width results in Figs. 5 and 6. Numbers of neurons contributing at each power are given in each panel; a neuron contributes at a given power only when a value is available both without illumination and at that power, so n differs between powers. Values of the index falling outside the interval 0–1, which arise from failed response fits, were excluded (four values per condition). P values are from two-tailed Wilcoxon signed-rank tests comparing each illumination power with the no-illumination baseline recorded in the same session; *P < 0.05, **P < 0.01, ***P < 0.001; n.s., not significant.

**Supplementary Table 1.** | Summary statistics for classical receptive field (CRF) data, related to Figs. 4 and 5. All values recomputed from the per-cell source workbooks. Each comparison is a two-tailed Wilcoxon signed-rank test on neurons with a value at both the no-illumination baseline and that power; means, s.d., median, s.e.m., and the paired baseline are over the same paired subset, so the baseline differs between powers by construction.

| Cohort | Measure | Power (mW mm<sup>−2</sup>) | n (paired) | Mean ± s.d. | Median | s.e.m. | P (unc.) | P (Holm) |
|---|---|---|---|---|---|---|---|---|
| PV-related ChR2 | HWHH (°) | 0 | 187 | 30.44 ± 13.55 | 26.60 | 0.99 | — | — |
|  |  | 160 | 145 | 27.99 ± 12.49 | 24.56 | 1.04 | 1.0 × 10<sup>−5</sup> | <1 × 10<sup>−4</sup> |
|  |  | 318 | 171 | 26.50 ± 11.86 | 23.60 | 0.91 | 6.4 × 10<sup>−6</sup> | <1 × 10<sup>−4</sup> |
|  |  | 477 | 177 | 23.86 ± 8.94 | 21.31 | 0.67 | 7.7 × 10<sup>−15</sup> | <1 × 10<sup>−4</sup> |
| PV-related ChR2 | Firing rate (spikes s<sup>−1</sup>) | 0 | 187 | 28.39 ± 28.06 | 19.61 | 2.05 | — | — |
|  |  | 160 | 145 | 26.27 ± 28.13 | 18.05 | 2.34 | 2.2 × 10<sup>−6</sup> | <1 × 10<sup>−4</sup> |
|  |  | 318 | 171 | 25.65 ± 26.39 | 18.38 | 2.02 | 1.5 × 10<sup>−12</sup> | <1 × 10<sup>−4</sup> |
|  |  | 477 | 177 | 24.41 ± 25.28 | 17.65 | 1.90 | 2.5 × 10<sup>−14</sup> | <1 × 10<sup>−4</sup> |
| PV-related ChR2 | Net firing rate (spikes s<sup>−1</sup>) | 0 | 187 | 22.64 ± 17.21 | 17.34 | 1.26 | — | — |
|  |  | 160 | 145 | 19.35 ± 15.83 | 14.62 | 1.32 | 2.5 × 10<sup>−7</sup> | <1 × 10<sup>−4</sup> |
|  |  | 318 | 171 | 19.79 ± 15.91 | 15.77 | 1.22 | 5.0 × 10<sup>−13</sup> | <1 × 10<sup>−4</sup> |
|  |  | 477 | 177 | 18.69 ± 15.15 | 14.97 | 1.14 | 2.1 × 10<sup>−15</sup> | <1 × 10<sup>−4</sup> |
| PV-related ChR2 | OSI | 0 | 187 | 0.36 ± 0.22 | 0.33 | 0.02 | — | — |
|  |  | 160 | 145 | 0.38 ± 0.22 | 0.37 | 0.02 | 0.069 | 0.208 |
|  |  | 318 | 171 | 0.39 ± 0.23 | 0.37 | 0.02 | 2.3 × 10<sup>−3</sup> | 9.3 × 10<sup>−3</sup> |
|  |  | 477 | 177 | 0.40 ± 0.24 | 0.39 | 0.02 | 1.2 × 10<sup>−5</sup> | 1 × 10<sup>−4</sup> |
| PV-related ArchT | HWHH (°) | 0 | 82 | 26.16 ± 9.54 | 25.34 | 1.05 | — | — |
|  |  | 160 | 82 | 25.85 ± 9.59 | 23.74 | 1.06 | 0.194 | 0.582 |
|  |  | 318 | 82 | 26.26 ± 9.24 | 23.78 | 1.02 | 0.959 | 1 |
|  |  | 477 | 82 | 26.96 ± 10.80 | 25.37 | 1.19 | 0.821 | 1 |
|  |  | 636 | 80 | 27.11 ± 9.53 | 26.59 | 1.07 | 0.124 | 0.497 |
| PV-related ArchT | Firing rate (spikes s<sup>−1</sup>) | 0 | 82 | 17.70 ± 9.91 | 15.78 | 1.09 | — | — |
|  |  | 160 | 82 | 18.86 ± 10.77 | 16.29 | 1.19 | 0.048 | 0.144 |
|  |  | 318 | 82 | 16.79 ± 8.66 | 14.67 | 0.96 | 0.0733 | 0.147 |
|  |  | 477 | 82 | 17.25 ± 10.90 | 14.14 | 1.20 | 0.153 | 0.153 |
|  |  | 636 | 82 | 16.40 ± 9.92 | 13.93 | 1.10 | 0.0041 | 0.0164 |
| PV-related ArchT | Net firing rate (spikes s<sup>−1</sup>) | 0 | 82 | 16.54 ± 9.45 | 15.06 | 1.04 | — | — |
|  |  | 160 | 82 | 17.96 ± 10.42 | 15.41 | 1.15 | 0.013 | 0.039 |
|  |  | 318 | 82 | 15.49 ± 8.04 | 13.75 | 0.89 | 0.0519 | 0.104 |
|  |  | 477 | 82 | 15.90 ± 10.51 | 13.46 | 1.16 | 0.0766 | 0.104 |
|  |  | 636 | 82 | 14.82 ± 9.28 | 12.71 | 1.02 | 3.7 × 10<sup>−4</sup> | 0.00149 |
| PV-only ChR2 | HWHH (°) | 0 | 50 | 34.15 ± 12.44 | 33.93 | 1.76 | — | — |
|  |  | 160 | 50 | 31.22 ± 10.84 | 30.92 | 1.53 | 0.079 | 0.315 |
|  |  | 318 | 47 | 31.24 ± 11.26 | 31.84 | 1.64 | 0.035 | 0.174 |
|  |  | 477 | 49 | 29.54 ± 11.88 | 29.53 | 1.70 | 9.1 × 10<sup>−4</sup> | 5.5 × 10<sup>−3</sup> |
| PV-only ChR2 | Firing rate (spikes s<sup>−1</sup>) | 0 | 54 | 12.91 ± 8.59 | 10.89 | 1.17 | — | — |
|  |  | 160 | 54 | 13.92 ± 9.69 | 10.81 | 1.32 | 0.40 | 0.971 |
|  |  | 318 | 54 | 14.88 ± 10.61 | 11.13 | 1.44 | 0.040 | 0.198 |
|  |  | 477 | 54 | 12.30 ± 8.43 | 10.77 | 1.15 | 0.32 | 0.971 |
| PV-only ChR2 | Net firing rate (spikes s<sup>−1</sup>) | 0 | 54 | 11.15 ± 7.60 | 9.45 | 1.03 | — | — |
|  |  | 160 | 54 | 12.41 ± 9.12 | 10.16 | 1.24 | 0.18 | 0.734 |
|  |  | 318 | 54 | 13.31 ± 9.95 | 10.11 | 1.35 | 8.2 × 10<sup>−3</sup> | 0.049 |
|  |  | 477 | 54 | 10.81 ± 7.24 | 8.89 | 0.99 | 0.56 | 1.0 |
| PV-only ChR2 | OSI | 0 | 133 | 0.46 ± 0.22 | 0.45 | 0.02 | — | — |
|  |  | 160 | 124 | 0.45 ± 0.22 | 0.45 | 0.02 | 0.363 | 0.740 |
|  |  | 318 | 125 | 0.45 ± 0.22 | 0.44 | 0.02 | 0.247 | 0.740 |
|  |  | 477 | 125 | 0.44 ± 0.22 | 0.44 | 0.02 | 0.022 | 0.089 |

**Supplementary Table 2.** | Summary statistics for full-field (FF) data, related to Figs. 4 and 5. Sources and conventions as in Supplementary Table 1.

| Cohort | Measure | Power (mW mm<sup>−2</sup>) | n (paired) | Mean ± s.d. | Median | s.e.m. | P (unc.) | P (Holm) |
|---|---|---|---|---|---|---|---|---|
| PV-related ChR2 | HWHH (°) | 0 | 205 | 24.13 ± 11.14 | 21.24 | 0.78 | — | — |
|  |  | 160 | 202 | 23.91 ± 10.81 | 21.02 | 0.76 | 0.99 | 1.0 |
|  |  | 318 | 198 | 23.87 ± 11.14 | 21.15 | 0.79 | 0.92 | 1.0 |
|  |  | 477 | 188 | 27.30 ± 17.53 | 22.38 | 1.28 | 0.095 | 0.285 |
| PV-related ChR2 | Firing rate (spikes s<sup>−1</sup>) | 0 | 205 | 27.03 ± 27.04 | 17.40 | 1.89 | — | — |
|  |  | 160 | 202 | 27.71 ± 27.20 | 19.12 | 1.91 | 4.8 × 10<sup>−3</sup> | 9.6 × 10<sup>−3</sup> |
|  |  | 318 | 198 | 27.48 ± 27.10 | 18.29 | 1.93 | 0.72 | 0.72 |
|  |  | 477 | 188 | 24.10 ± 26.03 | 15.33 | 1.90 | 6.3 × 10<sup>−5</sup> | 2 × 10<sup>−4</sup> |
| PV-related ChR2 | Net firing rate (spikes s<sup>−1</sup>) | 0 | 205 | 22.20 ± 18.23 | 16.28 | 1.27 | — | — |
|  |  | 160 | 202 | 22.99 ± 18.37 | 17.75 | 1.29 | 0.013 | 0.025 |
|  |  | 318 | 198 | 22.58 ± 18.73 | 16.52 | 1.33 | 0.80 | 0.80 |
|  |  | 477 | 188 | 18.77 ± 16.17 | 13.46 | 1.18 | 2.2 × 10<sup>−6</sup> | <1 × 10<sup>−4</sup> |
| PV-related ChR2 | OSI | 0 | 205 | 0.44 ± 0.25 | 0.41 | 0.02 | — | — |
|  |  | 160 | 202 | 0.45 ± 0.24 | 0.44 | 0.02 | 0.100 | 0.208 |
|  |  | 318 | 198 | 0.45 ± 0.25 | 0.46 | 0.02 | 0.90 | 0.90 |
|  |  | 477 | 188 | 0.37 ± 0.25 | 0.30 | 0.02 | 6.9 × 10<sup>−5</sup> | 3 × 10<sup>−4</sup> |
| PV-related ArchT | HWHH (°) | 0 | 82 | 19.20 ± 6.09 | 17.82 | 0.67 | — | — |
|  |  | 160 | 82 | 20.01 ± 6.66 | 19.52 | 0.74 | 0.285 | 0.285 |
|  |  | 318 | 82 | 21.05 ± 7.65 | 19.64 | 0.84 | 0.0401 | 0.12 |
|  |  | 477 | 82 | 22.03 ± 8.76 | 21.64 | 0.97 | 0.0078 | 0.0312 |
|  |  | 636 | 72 | 22.00 ± 12.14 | 19.40 | 1.43 | 0.0649 | 0.13 |
| PV-related ArchT | Firing rate (spikes s<sup>−1</sup>) | 0 | 82 | 9.78 ± 7.08 | 7.58 | 0.78 | — | — |
|  |  | 160 | 82 | 12.03 ± 8.74 | 9.41 | 0.96 | 3.3 × 10<sup>−6</sup> | 9.8 × 10<sup>−6</sup> |
|  |  | 318 | 82 | 12.80 ± 9.58 | 9.73 | 1.06 | 2.5 × 10<sup>−7</sup> | 1.0 × 10<sup>−6</sup> |
|  |  | 477 | 82 | 11.17 ± 7.66 | 8.80 | 0.85 | 0.0344 | 0.0344 |
|  |  | 636 | 82 | 11.25 ± 8.53 | 9.84 | 0.94 | 4.9 × 10<sup>−4</sup> | 9.8 × 10<sup>−4</sup> |
| PV-related ArchT | Net firing rate (spikes s<sup>−1</sup>) | 0 | 81 | 8.85 ± 6.73 | 6.75 | 0.75 | — | — |
|  |  | 160 | 81 | 11.05 ± 8.62 | 8.50 | 0.96 | 7.9 × 10<sup>−6</sup> | 2.4 × 10<sup>−5</sup> |
|  |  | 318 | 81 | 11.86 ± 9.27 | 8.72 | 1.03 | 2.6 × 10<sup>−7</sup> | 1.0 × 10<sup>−6</sup> |
|  |  | 477 | 81 | 10.10 ± 7.09 | 8.14 | 0.79 | 0.0603 | 0.0603 |
|  |  | 636 | 81 | 10.28 ± 7.90 | 9.10 | 0.88 | 5.7 × 10<sup>−4</sup> | 0.00115 |
| PV-only ChR2 | HWHH (°) | 0 | 125 | 29.97 ± 18.75 | 24.28 | 1.68 | — | — |
|  |  | 160 | 123 | 28.69 ± 18.96 | 23.61 | 1.71 | 0.30 | 0.888 |
|  |  | 318 | 121 | 32.83 ± 26.11 | 24.50 | 2.37 | 0.46 | 0.888 |
|  |  | 477 | 124 | 31.64 ± 23.32 | 23.99 | 2.09 | 0.33 | 0.888 |
| PV-only ChR2 | Firing rate (spikes s<sup>−1</sup>) | 0 | 128 | 9.83 ± 11.87 | 6.48 | 1.05 | — | — |
|  |  | 160 | 128 | 10.10 ± 13.11 | 6.53 | 1.16 | 0.74 | 0.971 |
|  |  | 318 | 128 | 9.15 ± 13.04 | 4.82 | 1.15 | 0.021 | 0.127 |
|  |  | 477 | 128 | 9.63 ± 13.06 | 5.63 | 1.15 | 0.20 | 0.813 |
| PV-only ChR2 | OSI | 0 | 129 | 0.55 ± 0.22 | 0.57 | 0.02 | — | — |
|  |  | 160 | 129 | 0.54 ± 0.24 | 0.55 | 0.02 | 0.264 | 0.740 |
|  |  | 318 | 129 | 0.49 ± 0.24 | 0.45 | 0.02 | 8.5 × 10<sup>−5</sup> | 5 × 10<sup>−4</sup> |
|  |  | 477 | 129 | 0.50 ± 0.26 | 0.49 | 0.02 | 5.8 × 10<sup>−3</sup> | 0.029 |

