## Supplementary Figures for "Parvalbumin interneuron circuit sharpens orientation tuning via the extraclassical surround in cat V1"

Supplementary data

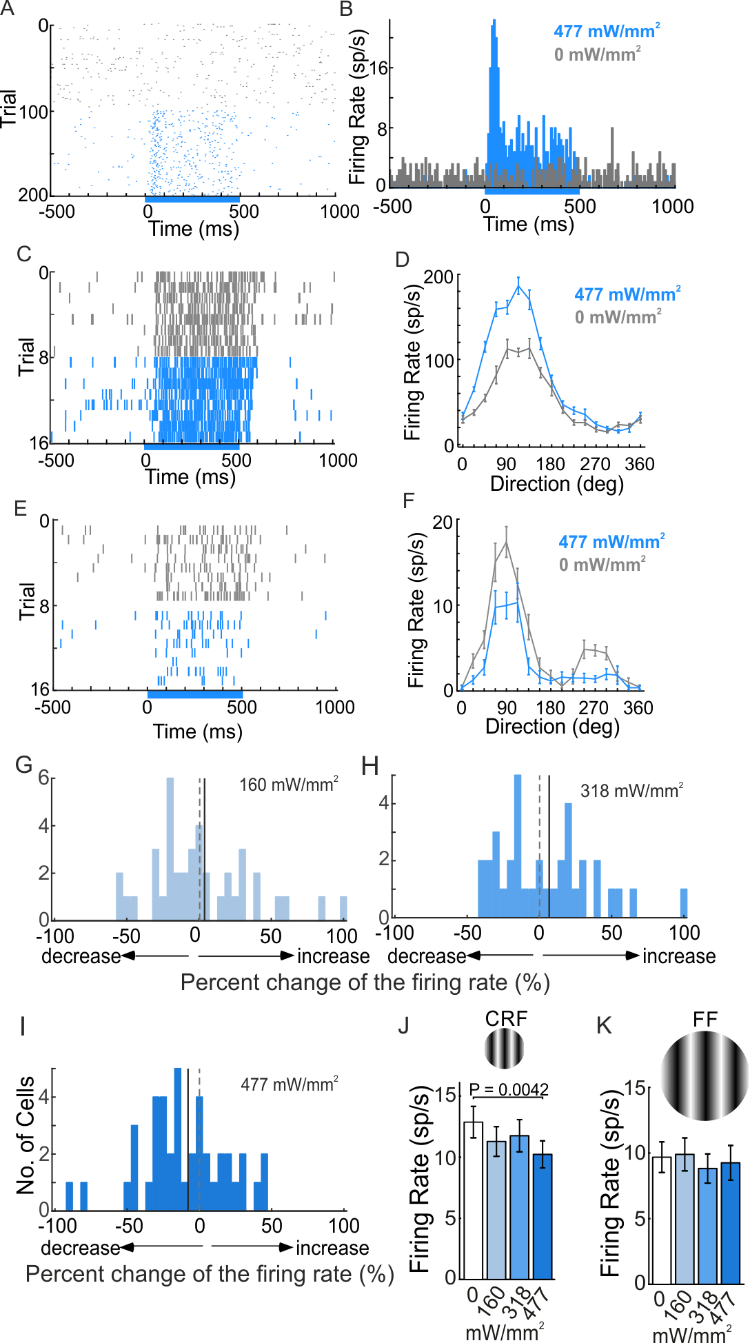

**Supplementary Fig. 1. Optogenetic activation of the PV-only circuit in vivo.** a, Raster of 100 trials of background activity (black) and 100 trials during 477 mW mm⁻² illumination (blue) for a ChR2-expressing neuron, presumed to be a PV interneuron. b, Peristimulus time histograms before (black) and during (blue) 477 mW mm⁻² illumination. c, Raster showing an increased response to drifting gratings during 477 mW mm⁻² blue illumination. d, Tuning curves for the neuron in c without illumination (black) and with illumination (blue). e, A neuron representative of the majority of recorded cells, showing a reduced visual response during illumination. f, Tuning curves for the neuron in e. g–i, Distributions of per-cell percentage change in firing rate at 160 (g), 318 (h) and 477 mW mm⁻² (i) for the PV-only circuit infected with the ChR2-YTx virus; solid lines mark the distribution mean and dashed lines zero. j, Population mean ± s.e.m. firing rate for CRF stimuli. k, Population mean ± s.e.m. firing rate for full-field stimuli. P values are from two-tailed Wilcoxon signed-rank tests comparing each illumination power with the no-illumination baseline.

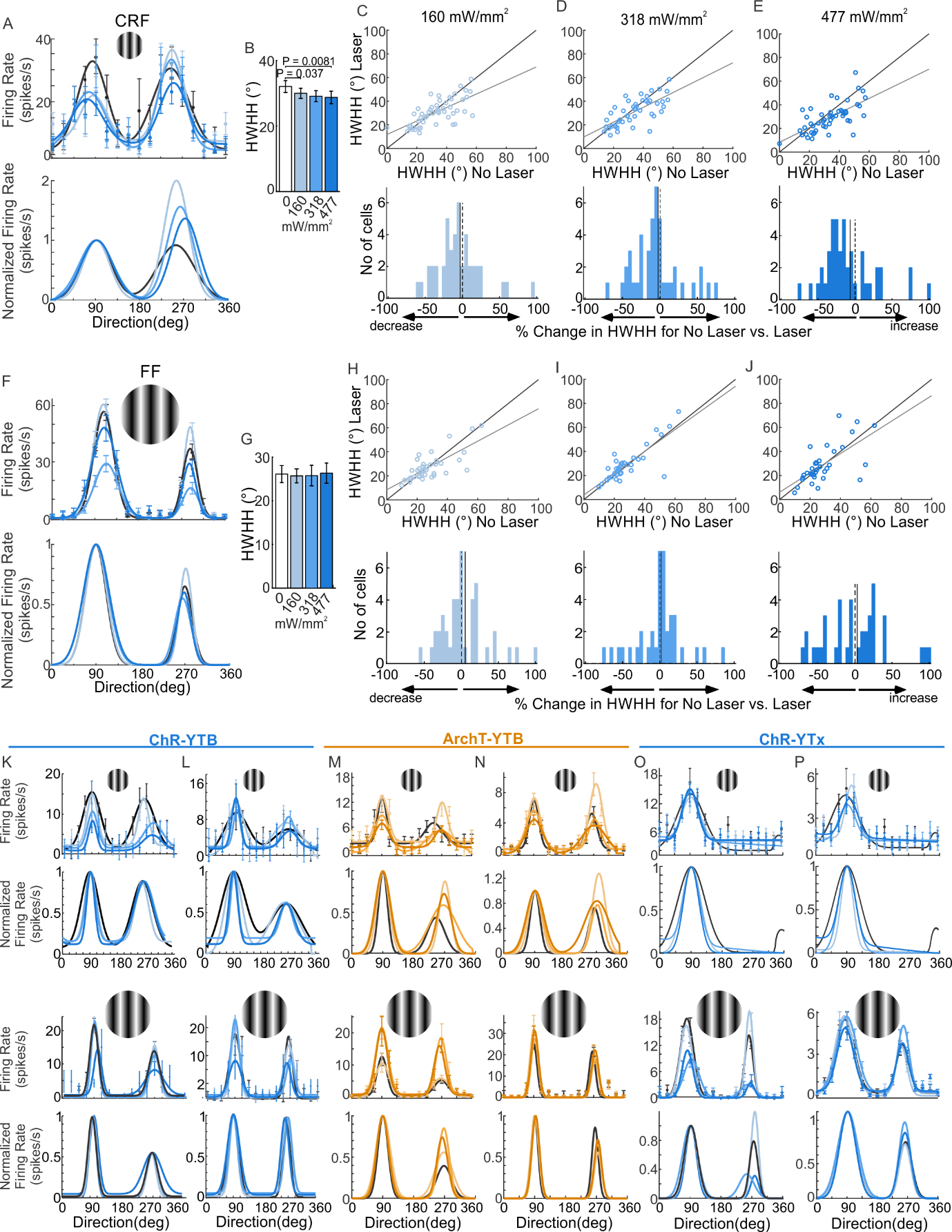

**Supplementary Fig. 2. Orientation tuning width in the PV-only circuit.** Half-width at half-height (HWHH) for the circuit infected with AAV-PV-YTx and EnvA-mCherry-ChR2-dG rabies virus, for classical receptive field (CRF) and full-field (FF) stimuli. a, Tuning curves from one neuron at each illumination power at the preferred stimulus size; top, mean responses with s.e.m. and Gaussian fits colour-coded by power; bottom, the same fits normalized. b, Mean CRF HWHH at each illumination power during ChR2 activation. c–e, Top, per-cell CRF HWHH at each of three illumination powers versus no illumination; bottom, distributions of percentage change from the no-illumination condition at 160, 318 and 477 mW mm⁻². Negative values indicate sharpening and positive values broadening. f, As in a, for full-field stimuli. g, Mean FF HWHH at each illumination power during ChR2 activation. h–j, As in c–e, for full-field stimuli; the grey solid line marks the distribution mean and the dashed line marks zero. k–p, Additional single-neuron examples. k,l, Top two rows, mean orientation tuning curves and fits for two neurons recorded at the optimal stimulus size without and with blue illumination (160, 318 and 477 mW mm⁻²) after AAV-PV-YTB and EnvA-mCherry-ChR2-dG injection, with normalized fits beneath; bottom two rows, the same for surround-engaging stimuli. m,n, As in k,l, after AAV-PV-YTB and EnvA-GFP-ArchT-dG injection. o,p, As in k,l, after AAV-PV-YTx and EnvA-mCherry-ChR2-dG injection. Data are mean ± s.e.m. P values are from two-tailed Wilcoxon signed-rank tests comparing each illumination power with the no-illumination baseline.

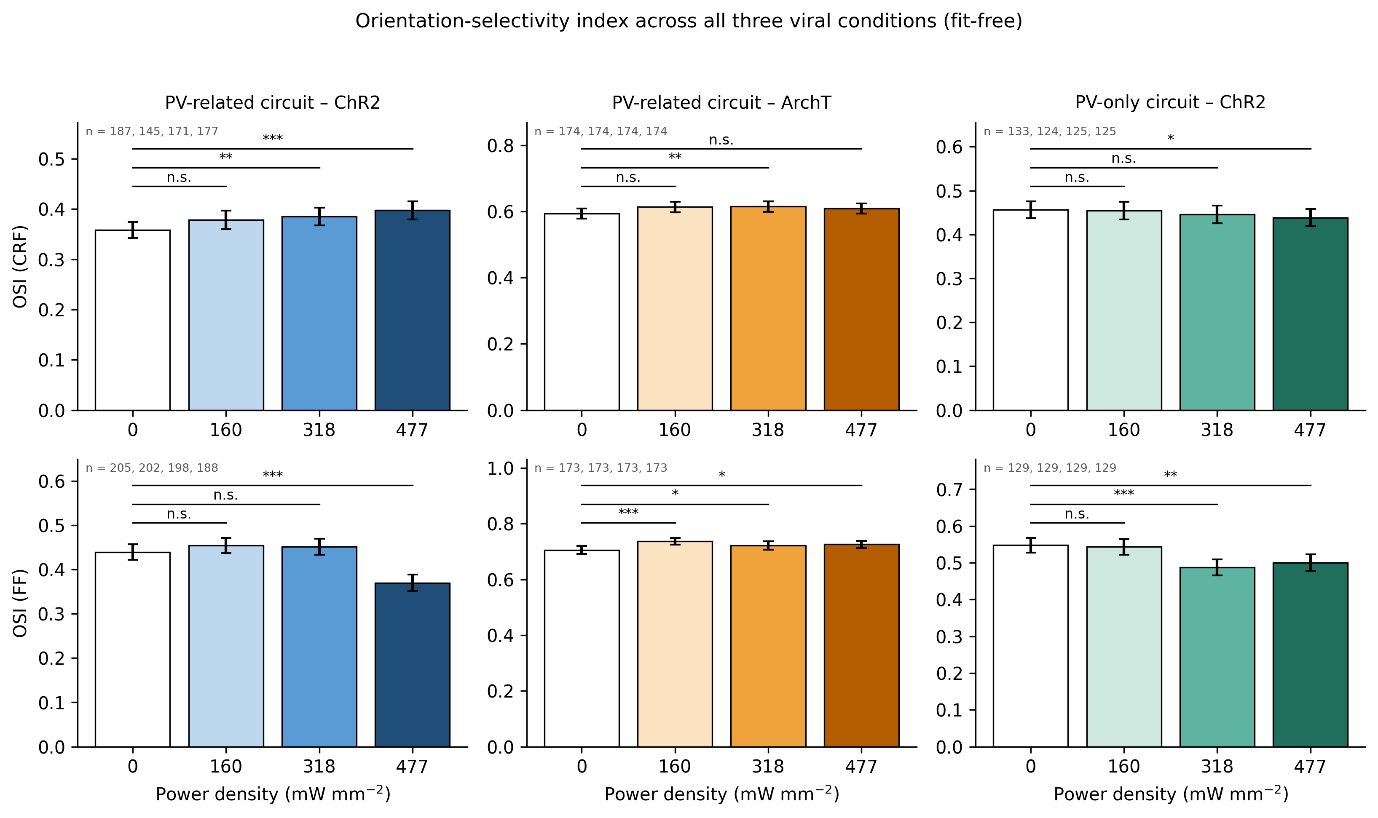

**Supplementary Fig. 3. Orientation-selectivity index across all three viral conditions.**

Mean orientation-selectivity index (OSI) ± s.e.m. against illumination power density for classical-receptive-field (CRF; top row) and full-field (FF; bottom row) stimuli, for the PV-related circuit expressing ChR2 (left), the PV-related circuit expressing ArchT (middle) and the PV-only circuit expressing ChR2 (right). The OSI is computed directly from the measured responses and does not depend on the Gaussian fit, and so provides a fit-free test of the tuning-width results in Figs. 5 and 6. Numbers of neurons contributing at each power are given in each panel; a neuron contributes at a given power only when a value is available both without illumination and at that power, so n differs between powers. Values of the index falling outside the interval 0–1, which arise from failed response fits, were excluded (four values per condition). P values are from two-tailed Wilcoxon signed-rank tests comparing each illumination power with the no-illumination baseline recorded in the same session; *P < 0.05, **P < 0.01, ***P < 0.001; n.s., not significant.
